# Multimodal immunopharmacologic screens identify drugs rewiring the cancer-immune interface

**DOI:** 10.64898/2026.04.13.717771

**Authors:** Jonas Bouhlal, Emmi Jokinen, Petra Nygren, Diogo Dias, Aleksandr Ianevski, Jay Klievink, Hanna Lähteenmäki, Salomé Decombis, Hanna Duàn, Emmi Järvelä, Emma Saarinen, Anna Näätänen, Konstantin Matjusinski, Tiina Kasanen, Tiina Hannunen, Laura Turunen, Mikko Myllymäki, Essi Laajala, Diana Schenkwein, Seppo Ylä-Herttuala, iCAN Flagship, Dean Lee, Matti Korhonen, Helka Göös, Tero Aittokallio, Maija Hollmén, Constantine Mitsiades, Sara Gandolfi, Olli Dufva, Satu Mustjoki

**Author notes:** Equal contribution (Emmi Jokinen, Petra Nygren). Equal contribution (Olli Dufva, Satu Mustjoki).

## Abstract

Natural killer (NK) cell-based therapies are a promising approach in cancer, but their efficacy is limited by impaired effector function and tumor-intrinsic resistance. To systematically identify therapeutic strategies that target both sides of the cancer-immune interface, we designed a multimodal immunopharmacologic screening platform comprising high-throughput co-culture drug screens, cytokine secretome profiling, single-cell perturbation screens, and genome-scale CRISPR screening, followed by validation in biobanked patient-derived models. Applying the platform across five blood cancer types, we identified protein kinase C (PKC) activation to simultaneously increase effector cytotoxicity and cytokine secretion through transcriptomic rewiring, and tumor susceptibility to NK cell killing through tumor-intrinsic PKC-δ. In patient samples, PKC activation sensitized NK-resistant leukemic progenitors to NK cell killing. In addition, NEDD8 inhibition enhanced NK function and shifted tumor TNF signaling towards pro-apoptotic pathways. Our platform provides a systematic approach to identify drugs rewiring both sides of the cancer-immune interface to circumvent tumor immune resistance.

## INTRODUCTION

Natural killer (NK) cells are central components of the innate immune system and play a critical role in the immune surveillance of malignant cells^1^. Unlike T cells, NK cells can recognize and eliminate malignant cells without prior sensitization, making them attractive candidates for cancer immunotherapy^2–4^. Several NK cell-based approaches, including adoptive transfer of expanded NK cells^2,5–7^, and genetically engineered CAR-NK cells^8,9^, are under clinical development, particularly in hematological malignancies such as acute myeloid leukemia (AML)^6,10,11^, multiple myeloma (MM)^12^, and lymphomas^8^. However, clinical responses have been variable, in part due to tumor resistance mechanisms and functional limitations of NK cells^13,14^.

NK cell cytotoxicity relies on the integration of activating and inhibitory receptor signaling^3,15^, cytoskeletal remodeling^16–18^, and the release of cytokines and lytic granules at the immunological synapse^18,19^. Multiple signaling modules - including the JAK-STAT axis downstream of γ-chain cytokines^20^, PI3K-AKT-mTOR networks that couple metabolic fitness to effector function^15,21,22^, canonical and non-canonical NF-κB signaling triggered by activating receptors^23–26^, and protein kinase C (PKC) -dependent checkpoints^27–29^ - have been established as key determinants of NK cell development and cytotoxic activity. In parallel, tumor-intrinsic processes such as metabolic rewiring^30,31^, defective death-receptor signaling^32^, and the secretion of immunosuppressive factors can further dampen NK cell responses^33–35^. Furthermore, recent genome-wide CRISPR screens in primary NK cells and target cells have identified novel genetic checkpoints and molecular vulnerabilities regulating NK cell cytotoxicity^36,37^. Despite these insights, how pharmacologic interventions can be optimally leveraged to simultaneously enhance NK cell activity and overcome tumor resistance remains to be defined. Small-molecule drugs can act on both immune and tumor cells, influencing their transcriptomic states, cytokine production by immune cells, or tumor-intrinsic signaling pathways that may not be captured at the transcriptomic level. Identifying drugs that target critical nodes in both immune and tumor cells and thereby enhance immune-mediated killing while preserving immune cell viability could enable the development of therapies that are inherently more resistant to escape, as resistance would require simultaneous alterations in two distinct processes. However, systematic approaches to identify such strategies are lacking.

Previous studies have primarily focused on individual compounds, pathways or disease types^38^, leaving a gap in systematic approaches to identify druggable mechanisms that enhance NK cell function across hematological cancers. Unbiased drug screening integrated with deep single-cell transcriptomic, cytokine profiling and CRISPR-based functional genomics offers an opportunity to uncover novel modulators of NK cytotoxicity and to dissect how pharmacologic agents reprogram both NK cells and target cells to favor immune-mediated killing. Such strategies could provide a foundation for rationally designed combination therapies that integrate NK cell-based immunotherapies with small-molecule drugs.

Here, we established an multimodal immunopharmacologic screening platform comprising sequential stages starting from high scale and moving towards increasingly detailed readouts. We first applied a high-throughput drug sensitivity and resistance testing (DSRT) platform to systematically evaluate the impact of more than 500 pharmacologic compounds on NK cell cytotoxicity against leukemia, lymphoma, and myeloma cell lines, while assessing the impact of these compounds on NK cell viability. Altogether, this approach generated over 13,000 dose-response curves, enabling a comprehensive assessment of how drugs modulate NK cell cytotoxicity across diverse contexts. Next, we selected the most potent compounds and generated single-cell perturbation and secretome atlases consisting of 240 conditions, allowing in-depth understanding of the effects at the cancer-immune interface. By integrating functional assays with single-cell transcriptomic profiling and secretome analysis of NK cells and target cells, we identified bryostatin 1, a PKC activator, and pevonedistat, an inhibitor of the NEDD8-activating enzyme (NAE), as two compounds that potently enhance NK cell-mediated killing. In the next stage, we systematically investigated the mechanisms of action of these highly promising compounds using co-culture genome-scale CRISPR screens and finally performed validations using biobanked clinically relevant patient samples. Mechanistic analyses revealed that these compounds act through distinct mechanisms, including activating receptor upregulation, cytoskeletal reprogramming, metabolic resilience, and sensitization of tumor cells to TNF-mediated apoptosis. Together, our findings uncover druggable pathways that reprogram NK cell-tumor cell interactions and highlight pharmacologic modulation as a promising avenue for next-generation NK cell-based therapies.

## RESULTS

### An immunopharmacologic screening platform identifies compounds modulating the cancer-immune interface

To systematically assess how drugs modulate NK cell cytotoxicity, we applied a high-throughput NK-tumor co-culture assay to profile 527 approved and investigational oncology compounds across 10 blood cancer cell lines (Figure 1A, S1A, Table S1), building on an assay framework described previously^39,40^. Cell line-specific effector-target (ET) ratios were optimized to achieve ∼50% NK-mediated target cell killing, enabling detection of both NK-enhancing and -suppressive drug effects (Figure S1B). We broadly examined NK-drug interactions across hematologic malignancies (AML, B-ALL, CML, MM, DLBCL), placing particular emphasis on AML due to its clinical urgency, substantial unmet therapeutic need, and the advanced clinical development of NK cell-based AML therapies.

**Figure 1:**
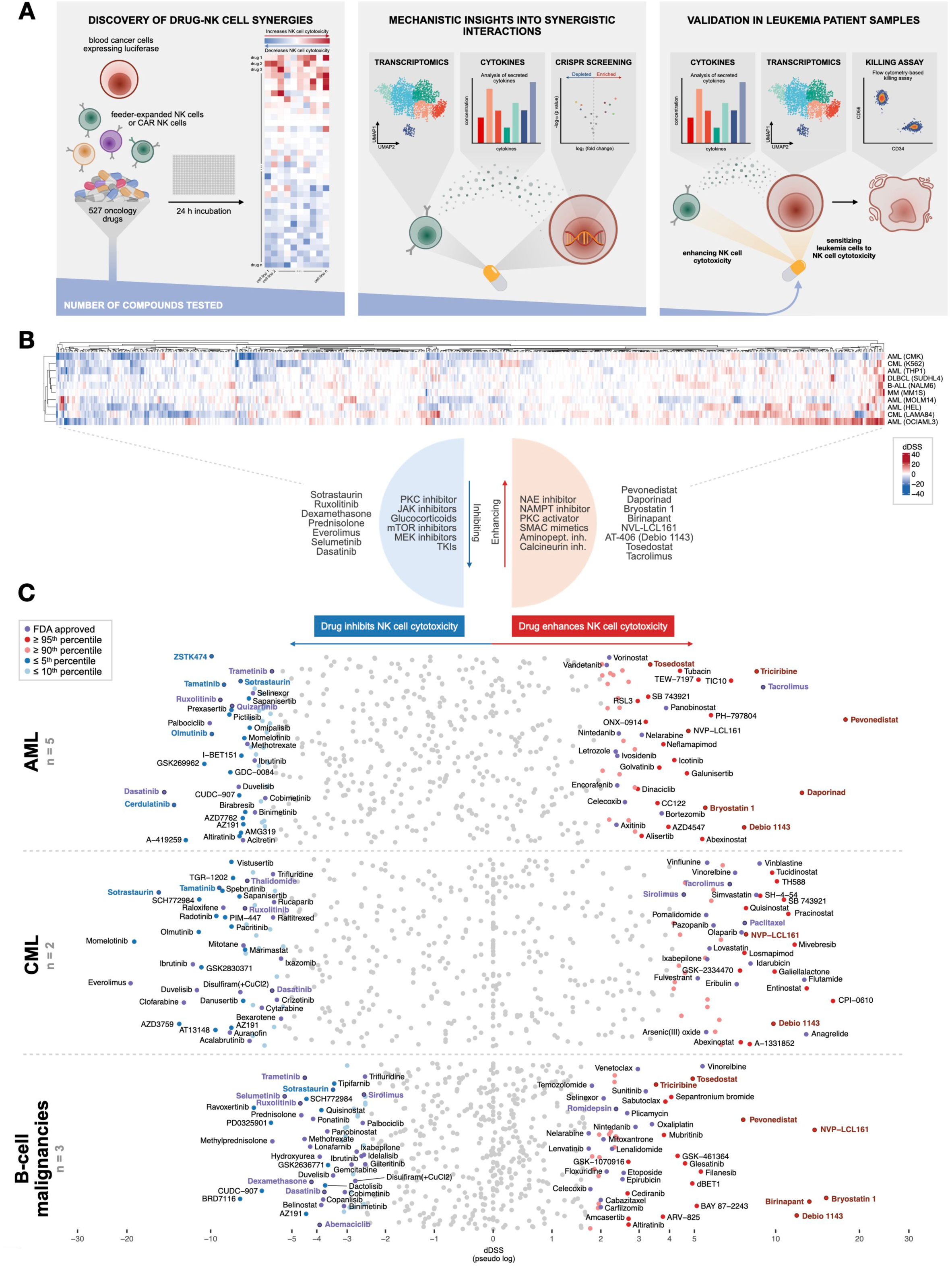
High-throughput drug resistance and sensitivity screening (DSRT) to identify compounds regulating NK cell cytotoxicity. (A) Overview of the workflow to identify NK cytotoxicity-modulating compounds and for deciphering underlying biological mechanisms. (B) Heatmap with overall results from DSRT experiments after 24h co-culture. Compounds increasing the NK cell-mediated killing of target cells appear red with a positive dDSS, whereas compounds decreasing the effectiveness of NK cells appear blue, indicating a negative dDSS. (C) Overview of how different compounds affect NK function in different blood cancers: Average pseudo-log transformed dDSS results from different cell lines, including 5 AML cell lines (MOLM-14, THP-1, OCI-AML-3, HEL, CMK), 2 CML cell lines (K562, LAMA-84), and 3 cell lines of B-cell malignancies (NALM-6, SU-DHL-4, MM1S). Compounds marked in bold were selected for further testing.

We assessed the effects of drugs on NK cell cytotoxicity by comparing the effect of the drug alone on target cells to that of drug+NK cell conditions and calculating the differential drug sensitivity score (dDSS). Overall, 77% of dDSS were within the range -5 to 5, indicating that most compounds had a limited effect on NK cell cytotoxicity, which could be observed as a minute difference between drug-treated and drug+NK-treated conditions. In contrast, some cell lines, such as OCI-AML-3, were substantially more responsive, exhibiting both stronger NK-enhancing and NK-inhibiting drug effects (Figure 1B, S1C). We observed disease-specific patterns, with concordant responses frequently shared among cell lines with a common cell of origin (Figure 1B and 1C). Across disease types, we identified compounds that either enhanced or inhibited NK cell cytotoxicity, reflecting their capacity to augment NK effector function, sensitize cancer cells to NK-mediated killing, or both (Figure 1C). Compounds enhancing NK cell cytotoxicity included pevonedistat (NAE inhibitor), daporinad (NAMPT inhibitor), bryostatin 1 (PKC activator), SMAC mimetics (birinapant, NVP-LCL161, AT-406), tosedostat (aminopeptidase inhibitor) and tacrolimus (calcineurin inhibitor). Conversely, sotrastaurin (PKC inhibitor), ruxolitinib (JAK inhibitor), glucocorticoids (dexamethasone, prednisolone), everolimus (mTOR inhibitor), selumetinib (MEK inhibitor) and dasatinib (tyrosine kinase inhibitor) showed NK cell inhibiting effects (Figure 1B and 1C).

### Drugs function consistently across diverse donors and potentiate CAR NK cell cytotoxicity

Consistent effects across donors and the ability to improve the function of CAR NK cells would be essential for clinical translation. To address this, we designed a custom drug panel based on our high-throughput DSRT results, containing compounds that consistently enhanced or inhibited NK cell cytotoxicity across all cell lines tested (average dDSS ≥ 6 or ≤ -6), as well as compounds with strong effects (dDSS ≥ 20 or ≤ -20) in at least one cell line. This panel was tested with NK cells from three healthy donors representing different HLA types to evaluate donor-dependence of drug effects and to validate our primary screening hits (Figure 2A, Table S2). We also assessed the applicability of these combinations in CAR NK cells by testing CD19-targeted CAR NK cells against NALM-6 in the presence of our custom drug panel. Furthermore, we assessed the effects of the compounds on NK cell viability using the CellTiter-Glo (GTC) assay.

**Figure 2:**
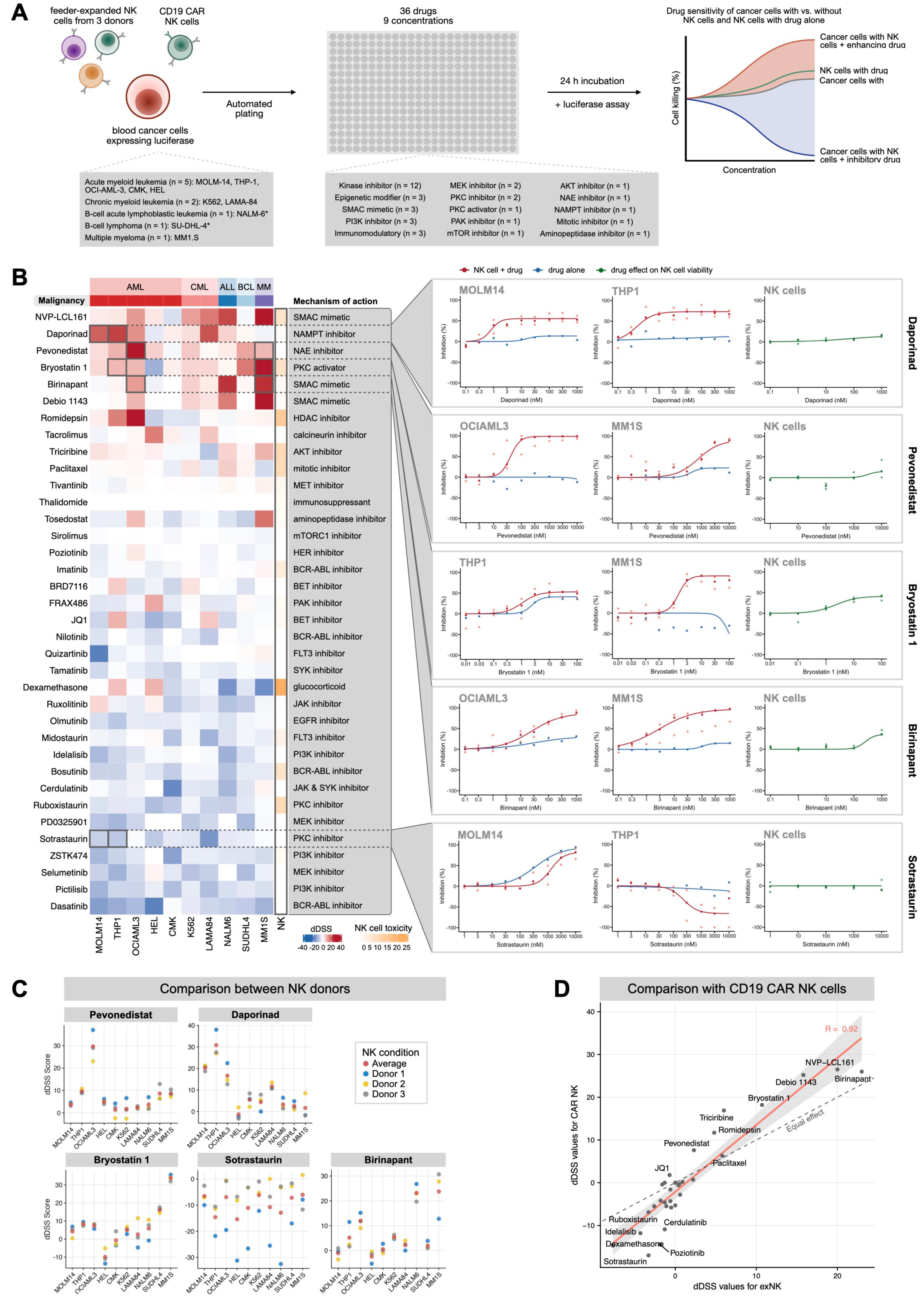
NK donor-dependent differences in drug effects and drug effects on CD19 CAR NK cells. (A) Validation experiment workflow with CD19 CAR NK cells and NK cells from three healthy donors representing different HLA-types. (B) Heatmap with average dDSS for each cell line from experiments with NK cells from three different donors combined with drugs selected for further study, in addition to DSS of NK cells alone to indicate the impact of drugs on their viability, with high scores indicating high toxicity to NK cells. To the right, inhibition curves of selected compounds. (C) Drug effects on NK cells from three different donors as a stacked dot plot of dDSS values. (D) Correlation of drug effects on non-CAR expanded NK cells (exNK) and CD19 CAR NK cells generated from the same donor when targeting NALM-6.

Validation screening confirmed our initial findings and highlighted compounds with consistent effects across multiple cell lines, independent of disease type (Figure 2B and S2A). Among the top enhancers, daporinad (NAMPT inhibitor) considerably increased NK cell cytotoxicity, particularly in AML cell lines such as MOLM-14 and THP-1, where the drug alone had little or no effect but, in combination with NK cells, induced up to 70% cancer cell killing without affecting NK cell viability (Figure 2B, S2A and S2B). Pevonedistat (NAE inhibitor) produced even stronger effects, especially in AML and MM cell lines. For example, in OCI-AML-3, pevonedistat alone was inactive but, in combination, achieved complete cancer cell eradication at ≥ 100 nM without reducing NK cell viability (Figure 2B, S2A and S2B). Bryostatin 1 (PKC activator) enhanced NK cell activity in MM, ALL, and AML, reaching up to 90% cancer cell killing in MM (Figure 2B and S2A). We identified 1 nM as an optimal concentration that maximized NK enhancement while minimizing NK toxicity observed at higher doses. Conversely, sotrastaurin (PKC inhibitor) produced broad and potent suppression of NK cytotoxicity across cell lines, consistent with its opposing mechanism to bryostatin 1 (Figure 2B and S2A). SMAC mimetics (NVP-LCL161, birinapant, debio 1143) also emerged as a notable drug class, potentiating NK cytotoxicity across diseases with minimal direct effects on cancer cells (Figure 2B and S2A), as observed previously in CML^40^.

The effects of most drugs were largely NK donor-independent, although the magnitude of enhancement or inhibition varied between donors. Notably, sotrastaurin’s inhibitory effect showed greater donor variability, potentially reflecting differences in PKC pathway expression or activity (Figure 2C).

Given the growing use of CAR-engineered NK cells in clinical trials^8,9^, we next examined whether drug-NK combinations could improve efficacy in this context. NK cells from the same donor, either unmodified feeder-expanded NK cells (exNK) or engineered with a CD19-CAR (CAR NK), were tested with the main hits from our panel. Cytotoxicity responses between CAR NK and exNK cells were strongly correlated (R = 0.92), indicating that the same compounds, which showed efficacy in the exNK setting, could also enhance and suppress CAR NK activity (Figure 2D and S2C). Importantly, several drugs exhibited even greater potentiating effects in CAR NK cells compared with unmodified NK cells, indicating that their ability to modulate the immune-cancer crosstalk is increasingly beneficial in the context of CAR NK cells.

### Single-cell transcriptomics and secretome profiling reveal drug-induced changes in NK cell function and target cell responses

To gain deeper insight into the biological mechanisms by which our top drug hits modulate NK cell-mediated cytotoxicity, we next examined their effects at the transcriptomic and proteomic levels (Figure 3A). Following the validation and prioritization of compounds with consistent and potent functional activity, we performed multiplexed single-cell RNA sequencing (scRNA-seq) on NK cells exposed to these drugs, either alone or in co-culture with cancer cells, resulting in a dataset consisting of 150,777 cells (33,195 NK cells and 117,582 cancer cells) and 240 conditions across 10 cell lines and 21 compounds (Figure 3B and S3A). Evaluating the single-cell perturbation atlas generated (Table S4B), NK cells showed distinct transcriptional profiles depending on the drug used and the target cell line with which they were in co-culture. Target cells also responded to NK cells and drugs, with some cell lines having more visible differences in the transcriptomic profile between NK-cultured and mono-cultured cancer cells (Figure S3A and Table S5). For example, NK-treated OCI-AML-3 cells clustered separate from those having been treated with drugs alone, compared to MM1S cells that had little difference between NK-treated and monoculture cancer cells, potentially reflecting their lower sensitivity to NK cells in general (Figure 3B and S3A). Drug-specific changes were also visible. For example, bryostatin 1-treated THP-1 cells appeared to have a similar transcriptomic profile as those having been co-cultured with NK cells, with an even more distinct separation in the THP-1 transcriptomic profile when treated with both NK cells and bryostatin 1 (Figure 3B and S3A).

**Figure 3:**
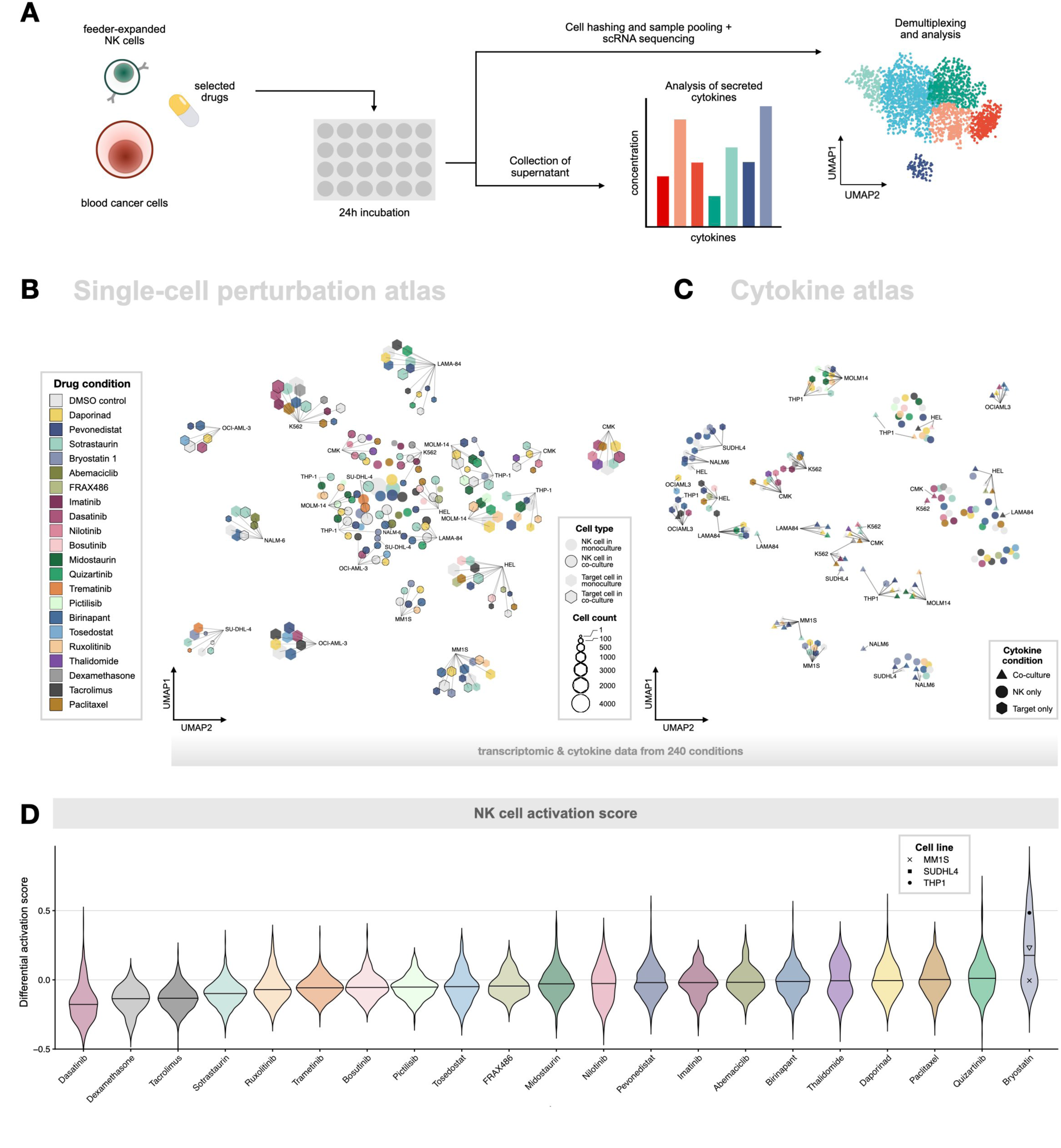
Single-cell transcriptomics and cytokine measurements from mono- and co-culture experiments with selected compounds. (A) Workflow of multiplexed single-cell RNA sequencing and analysis of secreted cytokines. (B) Overview of the 240 conditions from scRNA-seq experiments as pseudo-bulk UMAP, including mono- and co-cultured expanded NK cells and cancer cell lines, with and without drug treatment, shows that different drugs induce varying transcriptomic changes. Symbols without and with borders represent mono- and cocultured conditions, respectively. (C) Overview of cytokine secretion profiles from the 240 condition from scRNA-seq experiments highlights differences induced by different drug treatments and culture conditions. Visualized as UMAP of z-score-scaled cytokine levels. (D) Drug treatments’ effects on NK activation, visualized as violin plots of NK cell differential activation scores of drug-treated vs non-treated co-cultured NK cells. The scores were computed as the difference between mean activation scores from drug- and DMSO-treated cocultured NK cells based on previously published genes^32^ found upregulated in activated and type I IFN clusters, as illustrated in Figure S3C. Horizontal lines indicate median differential scores. For NK cells co-cultured in the presence of bryostatin 1, the target cell line used in each co-culture is indicated by symbols in the violin plot to highlight differences between responses to different target cell lines.

In parallel, culture supernatants from these experiments were analyzed to profile the spectrum of 27 secreted cytokines, generating 5184 cytokine measurements (Figure 3C). Similar to transcriptomic data, the levels of secreted cytokines reflected drug-induced changes. Aligned with observations made from scRNA-seq data, OCI-AML-3 cells alone with or without drug treatment had a largely different cytokine secretion profile compared to the conditions in which OCI-AML-3 cells were co-cultured with NK cells, reflecting the strong response by NK cells. Interestingly in THP-1, bryostatin 1 induced a cytokine profile that resembled that observed when co-cultured with NK cells without drug treatment. The cytokine profile of THP-1 and NK cells upon bryostatin 1 treatment was distinctly different from other THP-1 co-culture conditions, suggesting notable changes in the secretome and matching the observations made in the transcriptome (Figure 3C). Focusing on drug effects on NK cells’ cytokine secretion when cultured without target cells, we observed bryostatin 1 having similar, batch-independent effects (Figure S3B).

We further evaluated the effects of drugs on NK cell activation by calculating activation scores, comparing the expression of genes associated with activated and type I interferon-responsive NK cells^32^ in drug-treated NK cells co-cultured with cancer cells to that in DMSO-treated controls (Figure 3D, S3C and S3D). Similar to observations made from the large-scale transcriptomic and cytokine profiles, bryostatin 1 showed a notable increase in activation scores, especially in the case of THP-1, suggesting that PKC activation may improve NK cell responses to cancer cells.

This combined approach enabled us to link drug-induced transcriptional signatures with corresponding secretory responses, and thereby highlight perturbations that may explain effects observed in our DSRT screenings.

### Single-cell transcriptomics define drug-induced NK cell states

Building on insights from the transcriptomic and cytokine atlases, we next examined drug-induced transcriptomic changes in NK cells. Based on the gene expression profile, unsupervised clustering of expanded NK cells revealed nine clusters representing different functional states (Figure 4A). NK cells in monoculture with DMSO mostly belonged to the resting (cluster 1) and adaptive (cluster 8) clusters (Figure S4A and S4B). The resting cluster was defined by the expression of markers observed in expanded CD56 bright NK cells (*KLRC1*, *NCAM1*, *KLRB1*, *GZMA*, *GZMK*)^32^, and cells in the adaptive cluster based on the expression of *KLRC2*, *FCGR3A*, *GZMB*, *GZMH*, *LAG3* and *FCGR3A* (Figure 4A, S4C and Table S4A)^32^. Depending on the target cell in question, the co-culture with cancer cells induced two different cell clusters; activated (cluster 6) and type I interferon (cluster 5) (Figure 4A, S4A and S4B). Cells in the activated cluster expressed co-stimulatory (*TNFRSF4*, *TNFRSF9*, *CRTAM*) and inhibitory receptor genes (*HAVCR2*, *TIGIT*), genes related to death receptor ligands (*TNFSF10*, *ENTPD1*) consistent with previously reported findings (Figure 4A and S4C)^32^. In addition, NK cells in the activated cluster expressed *BCL2L11*, *PKM* and *RAMP1* (Figure 4A and S4C). Cells in the type I IFN cluster expressed genes related to interferon signaling activation and antiviral response (*IRF7*, *IRF9*, *IFIT1*, *IFIT3*, *IFI44L*, *ISG15*, *ISG20*)^32^, whereas NK cells belonging to the cytokine producing cluster expressed *CCL3*, *CCL4*, *TNF*, *IFNG* and *CD69* (Figure 4A and S4C).

**Figure 4:**
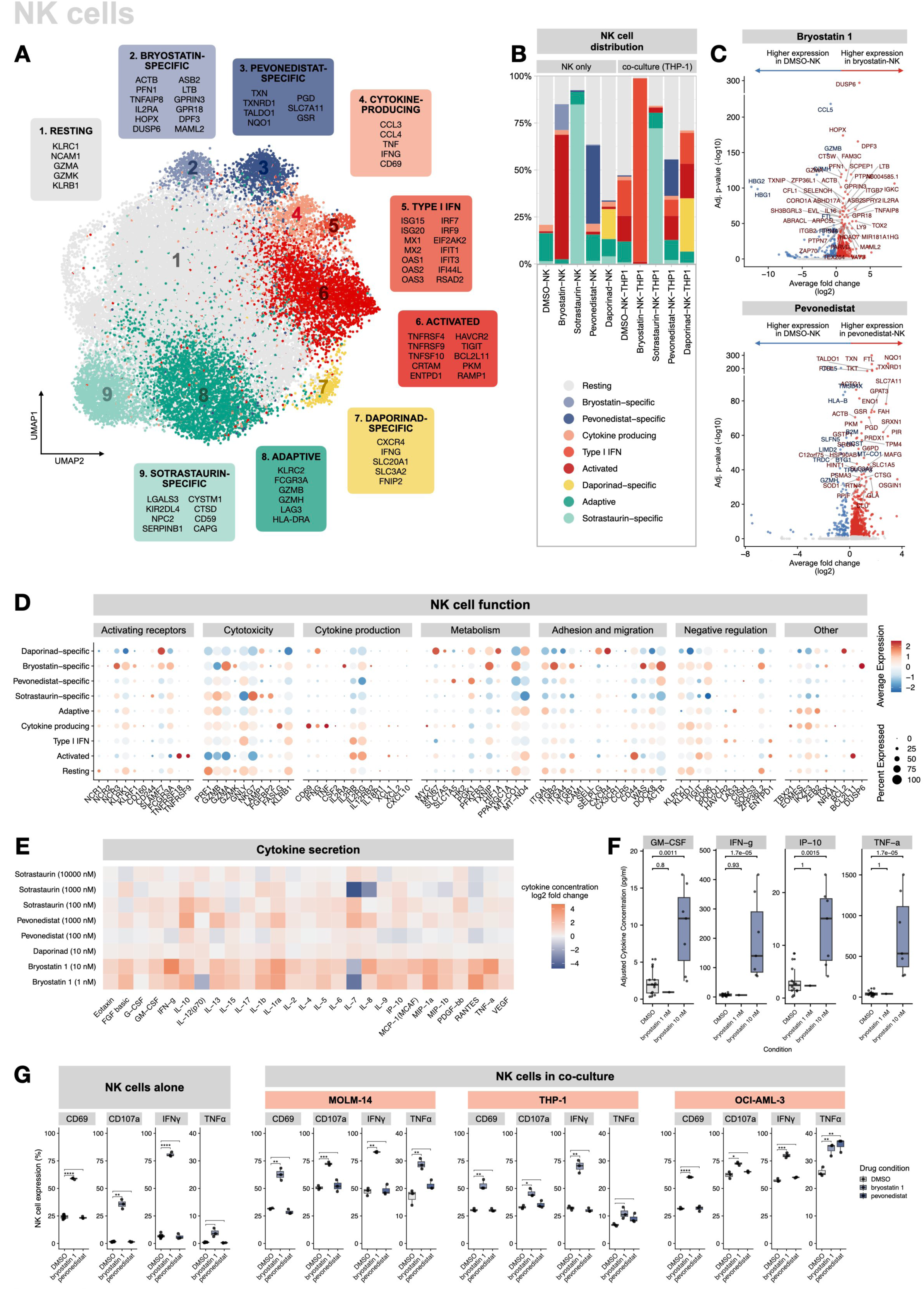
Evaluation of drug effects on NK cells by integrating data from scRNA-seq, cytokine and degranulation assays. (A) UMAP of all NK cells from mono- and co-culture conditions, including drug-treated NK cells. (B) Drug treatment-induced changes to NK cell phenotypes shown by a bar plot of percentages of NK cells per cluster by culture condition. (C) Bryostatin 1 (top) and pevonedistat (bottom) -induced transcriptomic changes to NK cells compared to controls, shown as volcano plots of differentially expressed genes. (D) Differences in gene expression affecting NK cell function between clusters illustrated as a dot plot of selected differentially expressed genes. (E) Drug treatment induced changes in cytokine secretion of monocultured NK cells, as a heatmap of mean fold changes between drug- and DMSO-treated NK cells. (F) Comparison of secreted cytokine concentration of GM-CSF, IFN-γ, IP-10 (CXCL-10 and TNF-α) between monocultured bryostatin 1 treated NK cells or controls via un-paired, two-sided Wilcoxon rank-sum tests for each cytokine. (G) Differences in NK cell activation and degranulation upon treatment with bryostatin 1 or pevonedistat and control, for both mono- and co-cultured NK cells. Statistical testing was performed with two-sided Student’s t-tests, separately for each marker within each condition, and resulting p-values were reported as significance levels (* < 0.05, ** < 0.01, *** < 0.001, **** < 0.0001, ns = not significant).

In addition to classical NK cell transcriptomic phenotypes present at baseline and cancer cell-induced activation states, we identified drug-induced NK cell states, which were present when NK cells were cultured with the drugs alone as well as in co-culture settings (Figure 4A, 4B, S4A). Out of all drugs tested (Table S3A), bryostatin 1, sotrastaurin, pevonedistat and daporinad induced drug-specific clusters in the NK cell transcriptome (Figure 4A, 4B, 4C, S4A and S4B).

### PKC controls the tumor-induced activation states in NK cells

The PKC activator bryostatin 1 induced notable increases in the proportion of activated (in co-culture with SU-DHL-4) and type I interferon (co-culture with THP-1) NK cells, when compared to controls (Figure 4B, S4A). More specifically, bryostatin 1 induced the upregulation of genes involved in cytoskeletal organization and cell structure (*ACTB*, *PFN1*), transcriptional regulation and chromatin remodeling (*DPF3*, *MAML2*, *HOPX*), signal transduction and immune activation (*IL2RA*, *DUSP6*, *GPR18*, *GPRIN3*), as well as apoptosis and immune modulation (*TNFAIP8*, *ASB2*, *LTB*) (Figure 4A, 4B, 4C, 4D, S4A, S4B, S4C). Furthermore, bryostatin 1 augmented NK cell function by upregulating the expression of activating receptors (*NCR3*, *KLRK1*, *KLRF1*, *CD244*, *SLAMF7*, *FCGR3A*) and genes related to adhesion and migration (*ITGAL*, *ITGB1*, *ITGB2*, *WAS*, *DOCK8*) (Figure 4D). In addition, bryostatin 1 influenced NK cell cytotoxic functions by upregulating the expression of *GZMA* and downregulating *GZMB* (Figure 4D). At the proteomic level, NK cells cultured alone, treated with 10 nM bryostatin 1, secreted higher levels of nearly all measured cytokines relative to controls across multiple experiments (Figure 4E). Similarly at a lower drug dose (1 nM), NK cells secreted higher levels of cytokines relative to controls, although effects with the lower dose were not significant when results were compared between experiments (Figure 4E and 4F). Notably, at 10 nM bryostatin 1 concentration, when inspecting cytokines classically considered relevant in the context of NK cells, GM-CSF (p = 0.0011), IFN-γ (p = 1.7e-05), IP-10 (p = 0.0015) and TNF-α (p = 1.7e-05) were significantly increased compared to NK cells cultured with the vehicle (DMSO) (Figure 4F).

In addition to transcriptome and secretome analyses, we measured NK cell degranulation when combined with bryostatin 1 alone and in co-culture with AML cell lines. Bryostatin 1 (10 nM) significantly increases the expression of the activation marker CD69 and degranulation marker CD107a already when given to NK cells alone, suggesting drug-induced pre-activation of the immune cells (Figure 4G and Table S7D). When in co-culture with AML cells, CD69 expression was significantly higher in bryostatin 1-treated NK cells compared to controls, similar as CD107a when in co-culture with MOLM-14 (Figure 4G and Table S7D). In addition, IFN-g production was significantly increased with bryostatin 1, both in NK cell monoculture and in co-culture with AML cell lines (Figure 4G and Table S7D).

The PKC inhibitor sotrastaurin, on the other hand, induced a sotrastaurin-specific cluster when using a 10,000 nM dose (Figure 4B, S4A). Cells in this cluster showed increased expression of inhibitory receptor-associated signaling (*KIR2DL4*), along with elevated *LGALS3* expression, lysosomal function and protease activity (*NPC2*, *CTSD*, *SERPINB1*), as well as genes related to membrane protection and cytoskeletal remodeling (*CD59*, *CAPG*, *CYSTM1*) (Figure 4A, S4C, S4D). Sotrastaurin also prevented NK cells from exhibiting classical activation signatures upon co-culture with cancer cells, in contrast to what was observed in NK cells having been in co-culture with the same cancer cells in control conditions (DMSO) (Figure 4B, S4A and S4B). A similar trend was observed in the case of tacrolimus, dasatinib and trametinib (Figure S3D, S4A and S4B). Instead, upon treatment with 10,000 nM sotrastaurin, most NK cells remained in the sotrastaurin-specific cluster (Figure 4B, S4A and S4B), whereas lower doses (100-1000 nM) of sotrastaurin allowed fewer NK cells to activate, with more NK cells to remaining in the resting cluster (Figure S4A and S4B). In contrast to the bryostatin 1 induced phenotype, sotrastaurin notably downregulated genes related to adhesion and migration (Figure 4D). Although clear changes were observed in the transcriptome, the analysis of secreted cytokines of NK cells cultured alone revealed less pronounced changes compared to bryostatin 1 (Figure 4E). Together, our findings identify PKC as a central node controlling the tumor-induced activation state in NK cells we previously identified, with NFAT, MAPK and Src pathways likely additionally contributing to inducing the activated state.

### NEDD8 and NAMPT inhibition reprogram genes controlling NK cell metabolism

Pevonedistat induced mostly changes downstream of NRF2, upregulating genes such as *TXN*, *TXNRD1*, *NQO1*, *TALDO, PGD* and *GSR* (Figure 4A, 4C, 4D, S4C and Table S4A, S4B). These transcriptional changes were observed both in mono- and co-culture conditions. In co-culture with AML cell lines, a subset of NK cells transitioned to the activated and type I IFN clusters, although the majority still remained in the pevonedistat-specific cluster (Figure 4B, S4A and S4B). Furthermore, pevonedistat induced a shift in metabolic regulation by upregulating genes related to amino acid transportation and glycolysis (*SLC1A5*, *SLC7A11, PGK1*) and resulted in the upregulation genes related to cytotoxic functions (*GZMB*, *GNLY*) (Figure 4A, 4C, 4D and S4C).

Daporinad-treated NK cells exhibited changes in metabolic regulation, with increases in the expression of genes related to glycolysis (*PGK1*) and amino acid transportation (*SLC3A2*, *SLC20A1*, *SLC7A5*) (Figure 4A, 4C, 4D and S4C). Daporinad also caused the upregulation of the proliferation marker *MKI67* and showed increases in *HIF1A* expression often observed upon hypoxic conditions (Figure 4D). Strikingly, daporinad caused the upregulation of *IFNG* and *CXCR4* expression, potentially explaining partly the increased cytotoxic capabilities in AML cell lines (Figure 4A, 4C, 4D and S4C).

### PKC activation reprograms cancer cells to a more NK cell-susceptible state

In addition to characterizing NK cell responses, our scRNA-seq and cytokine profiling data also provided insights into drug-induced changes in cancer cells, including priming them for increased susceptibility to NK cell-mediated killing. We focused on PKC pathway modulators, due to the strong enhancing effects of bryostatin 1 in AML and myeloma cell lines, as well as the strong inhibiting effects of sotrastaurin across different malignancies. Bryostatin 1, a macrocyclic lactone binding to the C1 domain of PKC, activates the enzyme and has been shown to reduce tumor cell proliferation and promote apoptosis in numerous hematological and solid tumor cell lines (Figure 5A)^41–44^. However, its benefit as a single agent has remained minimal^45^, in accordance with our DSRT results (Figure 2B and S2A). Sotrastaurin, a pan-PKC inhibitor, on the other hand, reduces the activity of different isoforms of protein kinase C (Figure 5A).

**Figure 5:**
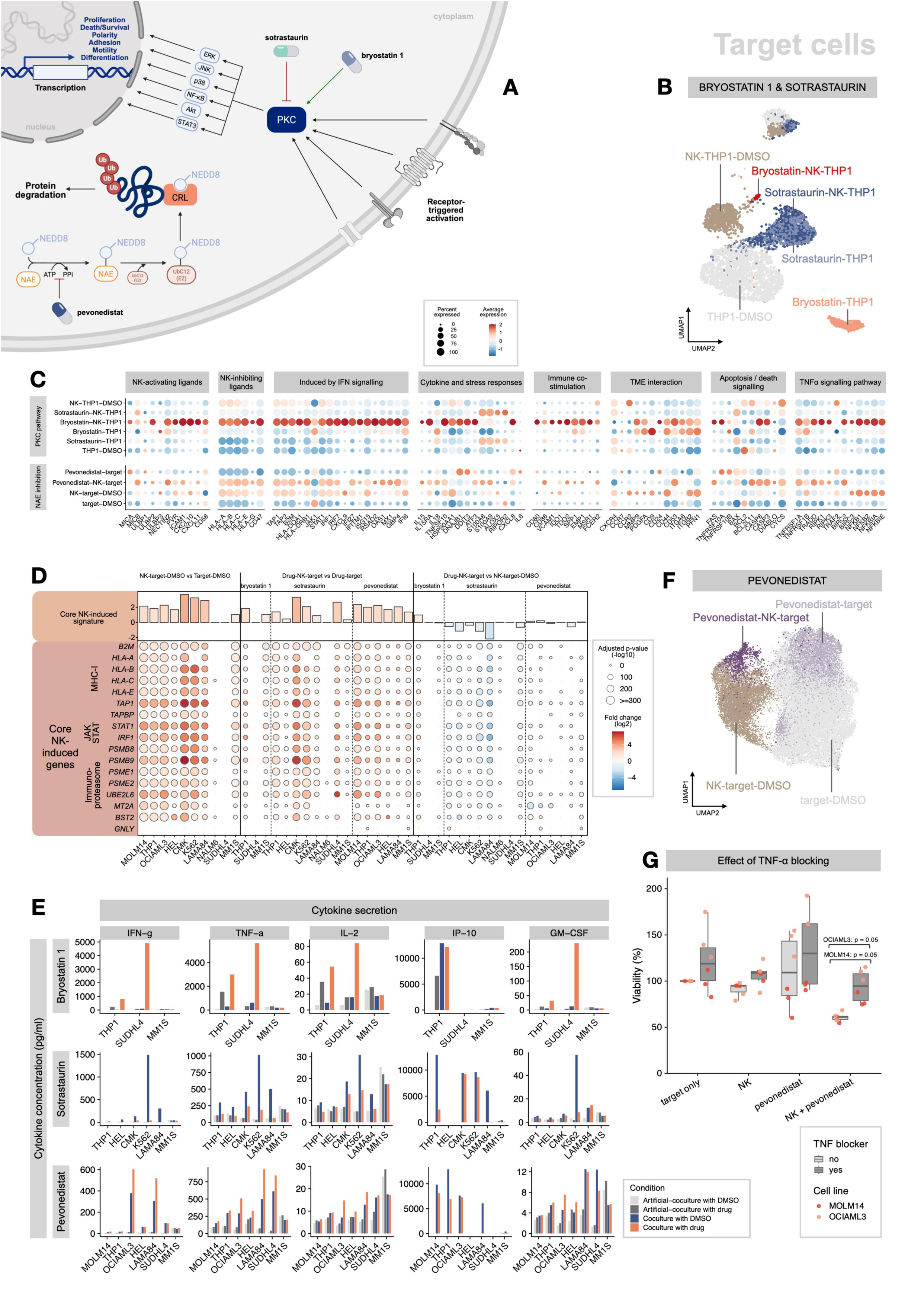
Drug and NK cell effects reflected on target cells transcriptomic and cytokine profile. (A) Illustration of mechanisms of action of PKC modulators and NEDD8 inhibitors. Adapted from Kazanietz et al.^71^ and Fathi^72^. (B) UMAP of target cells (THP-1) reflecting different transcriptional profiles depending on treatment with PKC modulators and/or NK cells. (C) Dot plot of selected, differentially expressed genes in target cells reflecting interactions with NK cells and changes depending on drug treatment. Gene expression profiles of PKC modulators (top) shown in THP-1, NAE inhibition with pevonedistat (bottom) includes all target cells having been treated with pevonedistat (MOLM-14, THP-1, OCI-AML-3, HEL, LAMA-84, MM1S) with and without NK cells. (D) Differences in core NK-induced genes between conditions in the titles as a dot plot of fold changes. Bar plots on the top illustrate the average fold changes over all the genes. (E) Bar plots of cytokine concentrations measured from supernatant. Artificial co-culture includes the sum of target and NK cell mono-culture cytokine concentrations. (F) UMAP of target cells (MOLM-14, THP-1, OCI-AML-3, HEL, LAMA-84, MM1S) treated with pevonedistat and controls. (G) Effect of TNF-⍺ blocker infliximab on MOLM-14 and OCI-AML-3 cells in combination with pevonedistat and/or NK cells. Statistical significance tested separately for both cell lines using un-paired, one-sided Wilcoxon rank-sum test.

Bryostatin 1 reshaped the cancer cell transcriptome both alone and in NK co-culture, shifting cells into a distinct state while still aligning them near NK-only-treated cells (Figure 5B, 5C and S5A). The analysis of NK-induced gene signatures^32^ further supported this finding, revealing that bryostatin 1-treated cancer cells in NK cell co-culture present higher expression of core NK-induced genes when compared to DMSO controls (Figure 5D). These findings suggest that bryostatin 1 primes target cells toward an NK-responsive phenotype yet drives its own transcriptional program.

In contrast, sotrastaurin held THP-1 cells in a shared cluster regardless of NK cell exposure. Drug-treated cells failed to acquire the transcriptional changes normally triggered by NK co-culture, reflecting that sotrastaurin (10,000 nM) prevents target cells from engaging typical NK-induced responses due to blocked NK cell activation (Figure 5B, 5C, 5D and S5A). However, this effect was not observed as clearly in lower dose of 1000 nM (K562) and only moderately with 100 nM (CMK) (Figure 5D and S5A).

Bryostatin 1 alone upregulated the NK-activating ligands *ULBP3*, *NECTIN2* and *CD58* in cancer cells, in addition to upregulating the expression of genes related to tumor microenvironment (TME) interaction and differentiation (*CD9*, *CD24*, *CD44*, *CD53*, *MMP9*, *PDGFA*, *ITGA6, ITGB2*) (Figure 5C and Table S5). Moreover, the expression of genes related to interferon signaling were upregulated, including *JAK1*, *ISG15*, *OAS1*, *MX1*, and *IFI6*. In addition, genes related to the TNF-α signaling pathway and death receptor signaling, such as *TNFRSF10B*, *BCL2L11*, *TNFRSR1A* and *TNFRSR1B*, were upregulated. Overall, these findings suggest that PKC activation primes cancer cells to a more NK-stimulating and apoptosis-susceptible state.

Cells treated with both bryostatin 1 and NK cells exhibited strong upregulation of genes involved in multiple pathways that underlie NK cell activation and cytotoxic function against cancer cells (Figure 5C). These included the upregulation of genes related to both activating (*MICA*, *MICB*, *ULBPP2*, *PVR*, *ICAM1*, *CXCL10*, *CXCL11*, *CD58*) and inhibiting ligands (*HLA-A/B/C/E/G*, *CD47*), those induced by interferon signaling (*TAP1/2*, *B2M*, *STAT1*, *IRF1*, *IRF7*, *IFI27*, *IFITM1*, *ISG15*, *OAS1*, *MX1*), cytokine and stress responses (*IL15*, *IL15RA*, *TNFSF10*, *DNAJB1*, *DDIT3*, *ATF3*), immune modulation (*CD80*, *CD86*, *VCAM1*, *IDO1*, *TDO2*, *SPP1*, *EMP1*, *MSR1*, *FCER2*) as well as the TNF-α signaling pathway (*TNFRSF1A*, *TNFRSF1B*, *TRADD*, *RIPK1/3*, *BIRC2/3*, *NFKB1/2/IA/IE*) and death signaling (*FAS*, *TNFSFSRF10A*, *CASP8*, *CASP3*) (Figure 5C).

In the case of sotrastaurin, these genes were largely downregulated across the board, indicating the inactivity of these pathways and mechanisms identified as important in the context of PKC activation. Sotrastaurin alone on cancer cells caused minimal impact on gene expression signatures related to NK cell interaction (Figure 5C). In line with reduced NK cell activation observed with sotrastaurin treatment, co-culture with NK cells in combination with sotrastaurin caused notably milder responses compared to controls, with lower NK cell-inhibiting ligand expression, and signatures corresponding to interferon, TNF-α and death receptor signaling (Figure 5C).

The cytokine secretion profiles matching scRNA-seq experiments revealed similarly striking patterns. We calculated cytokine concentrations for artificial co-cultures by combining cytokine concentrations from target-only and NK cell-only conditions and compared those to the concentrations measured from true co-cultures, with the aim to identify the additional cytokine secretion caused by NK-target interactions. In THP-1 and SU-DHL-4 co-cultures, bryostatin 1 treatment led to marked increases in IFN-γ, TNF-α, IL-2, and GM-CSF secretion, a response absent in the corresponding control conditions (Figure 5E). Interestingly, this effect was not observed in the case of MM1S, possibly due to a lower concentration (1 nM instead of 10 nM). In contrast, sotrastaurin caused these cytokines to be notably less secreted upon co-culture across multiple cell lines, indicating impaired secretion due to PKC inhibition (Figure 5E). These results suggest that PKC modulation has strong effects on NK cells’ secretory machinery.

### NEDD8 inhibition promotes TNF-dependent cancer cell apoptosis

We also further explored the mechanisms behind NAE inhibitor pevonedistat, which functions by inhibiting NAE, preventing it from activating cullin-RING ligases, resulting in impaired proteasome-mediated protein degradation and leading to apoptosis in cancer cells (Figure 5A). Similar to the transcriptomic data from NK cells, pevonedistat caused upregulation of genes downstream of NRF2 in target cells, which can be explained by its mechanism of action, leading to the accumulation of NRF2 (Table S5).

NAE inhibition prominently altered TNF-α signaling, affecting death-receptor pathways and apoptosis. Target cells exposed to both pevonedistat and NK cells showed marked upregulation of the TNF-receptor genes *TNFRSF1A* (TNFR1) and *TNFRSF1B* (TNFR2), and several downstream mediators of TNF-receptor signaling - including *RIPK1*, *RIPK3*, *BIRC2*, *CASP3*, *CASP4*, and *CASP8* - were similarly upregulated (Figure 5C). Notably, these expression levels differed substantially from those in cells treated with pevonedistat or NK cells alone, indicating a distinct transcriptional program induced by the combination (Figure 5C and 5F). In contrast to co-cultures with NK-only, NF-κB-related genes (*NFKB1*, *NFKB2*, *NFKBIA*, *NFKBIE*) were expressed at low levels in the combination treatment, potentially indicating a shift to enhanced pro-apoptotic effects. Moreover, interferon-stimulated genes (*OAS1*, *MX1*, *GBP4*) were slightly more upregulated in cells treated with pevonedistat and NK cells compared with NK cells alone (Figure 5C). Pevonedistat induced marginal effects to core NK-induced gene expression, suggesting that the increased killing resulted in by combining pevonedistat with NK cells is caused by alternative signaling pathways to those induced by NK cells alone (Figure 5D).

In addition to effects on TNF-α signaling on transcriptomic level, TNF-α was increasingly secreted in co-cultures where pevonedistat was applied across all cell lines tested (Figure 5E). To test our hypothesis that TNF-α signaling plays a key role in the pevonedistat-NK cell synergy, we added the TNF-α -blocking antibody infliximab to the co-culture assays of two AML cell lines (MOLM-14 and OCI-AML-3). The results supported this mechanism: in the presence of infliximab, the viability of leukemic cells in NK cell + pevonedistat conditions increased from approximately 50% to nearly 100% (p = 0.05), effectively abolishing the enhanced cytotoxic effect observed with pevonedistat in combination with NK cells (Figure 5G).

### CRISPR screening reveals cancer cell-intrinsic drivers of PKC and NEDD8-mediated immune sensitization

Building on the insights gained from single-cell RNA sequencing and cytokine profiling, we next sought to further dissect underlying tumor-intrinsic factors influencing sensitivity to the combination of drugs and NK cells through genome-wide CRISPR screening. We selected an AML cell line (THP-1) in which both bryostatin 1 and pevonedistat enhanced NK cell cytotoxicity and transduced it with the Brunello genome-wide CRISPR knockout (KO) library (Figure 6A).

**Figure 6:**
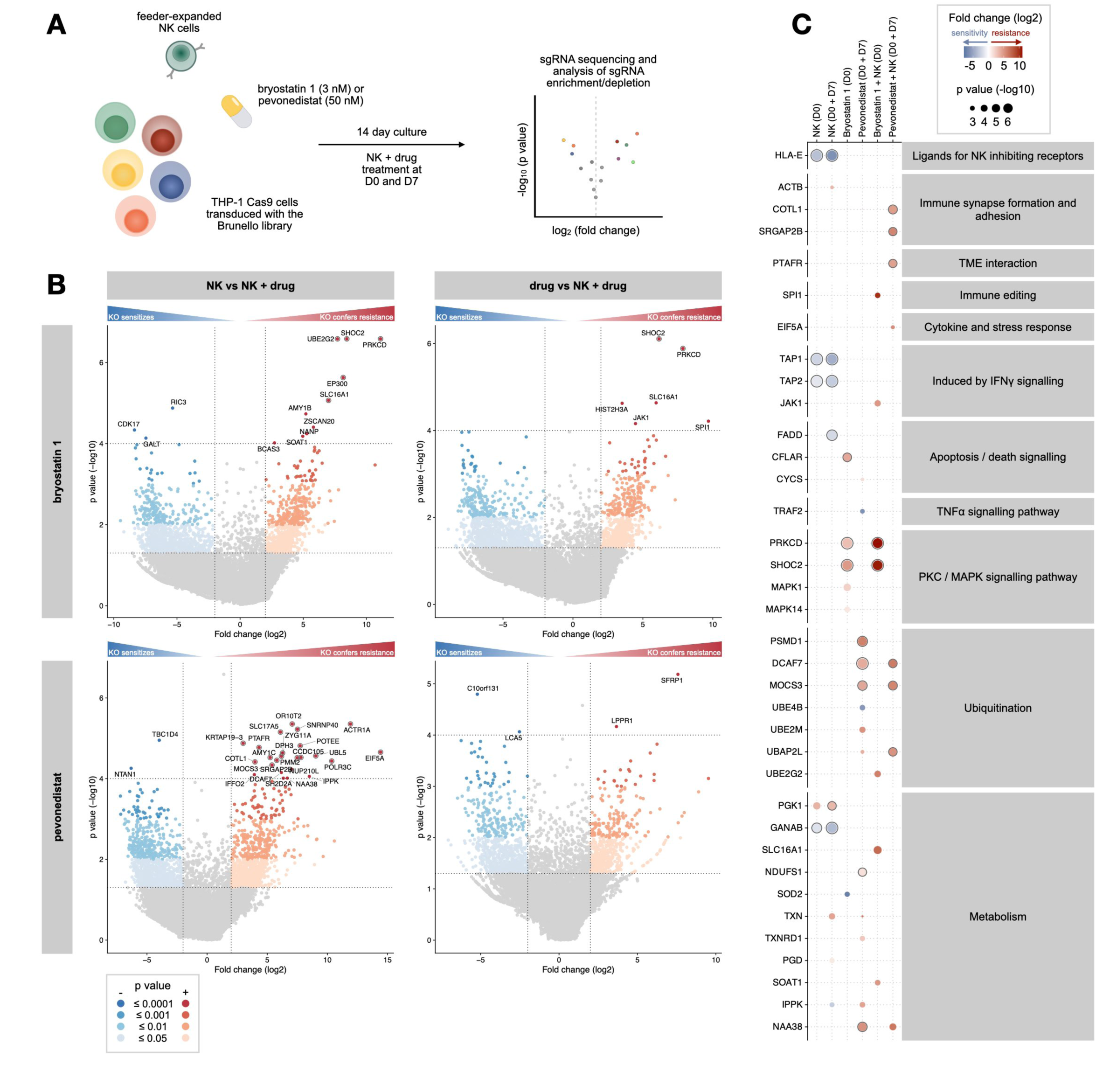
Genome-wide CRISPR screens of target cell resistance and sensitivity mechanisms to NK cell-drug combinations. (A) Workflow of genome-scale CRISPR screens. (B) Volcano plots of CRISPR results highlighting gene perturbations driving resistance or sensitivity to the combination of NK cells and bryostatin 1 (top) or pevonedistat (bottom). NK+drug-treated conditions are compared to NK-only-treated (left) and drug-treated (right) to identify gene perturbations specific to NK-drug combinations. Gray circles around gene points indicate a false discovery rate (fdr) ≤ 0.05. (C) Dot plot of gene perturbations, with selected genes relevant in the context of NK cells with p-value ≤ 0.05, log2 fold change ≥ 1, fdr ≤ 0.3. Dark circle indicates fdr ≤ 0.05.

In the baseline comparison of untreated THP-1 cells and NK-treated counterparts after 14 days of culture, cells with KO of *HLA-E*, *NLRC5*, *TAP1*, *TAP2*, *TAPBP* and *GANAB* were markedly depleted (Figure S6A and Table S6), confirming that the loss of these genes sensitizes target cells to NK cell-mediated killing, consistent with previous reports^32^, confirming the accuracy and biological relevance of our screening approach.

*PRKCD* (PKC-δ), *SHOC2*, *SLC16A1*, and *JAK1* knockouts were significantly enriched under selection with bryostatin 1 (3 nM) in combination with expanded NK cells (Figure 6B). KO of *PRKCD* and *SHOC2* in the context of bryostatin 1, led to resistance already in the drug-only treated cells (Figure 6C). However, the effect was further pronounced when combined with NK cells, suggesting that bryostatin 1 could sensitize cancer cells to NK cell killing via *PRKCD* and *SHOC2* mediated pathways and that cancer-intrinsic PKC activation contributes to the efficacy of the drug (Figure 6C). Only a few of the genes traditionally observed to be involved in causing resistance to NK cells, such as *HLA-E*, appeared as hits, indicating that treatment with a PKC agonist may circumvent MHC class-mediated resistance towards NK cells (Figure S1A and 6C).

For pevonedistat, the KO of *DCAF7*, *MOCS3* and *UBAP2L*, which play a role in protein ubiquitination, resulted in resistance to the drug alone and also to its combined effects with NK cells (Figure 6B and 6C). Genes of which the KO induced resistance specifically to the combined treatment with pevonedistat and NK cells included *COTL1*, *SRGAP2B* and *PTAFR* (Figure 6B and 6C). Together, the enrichment of these KO populations suggests that cytoskeletal regulation and stress/inflammatory signaling pathways may modulate tumor-cell vulnerability to pevonedistat-augmented NK cell killing.

### PKC activation overcomes immune resistance of patient-derived leukemic progenitors

Finally, we sought to evaluate the applicability of our drug-NK cell combinations in primary leukemia samples, moving a step closer to clinical translation. We selected bone marrow samples from eight AML and two B-ALL patients. Using a flow cytometry-based killing assay, we were able to identify the effects of NK cells and drugs, both alone and in combination, on primary bone marrow samples, while also performing scRNA-seq on the AML + NK cell co-cultures as well as studying effects on the secreted cytokines (Figure 7A, S7A).

**Figure 7:**
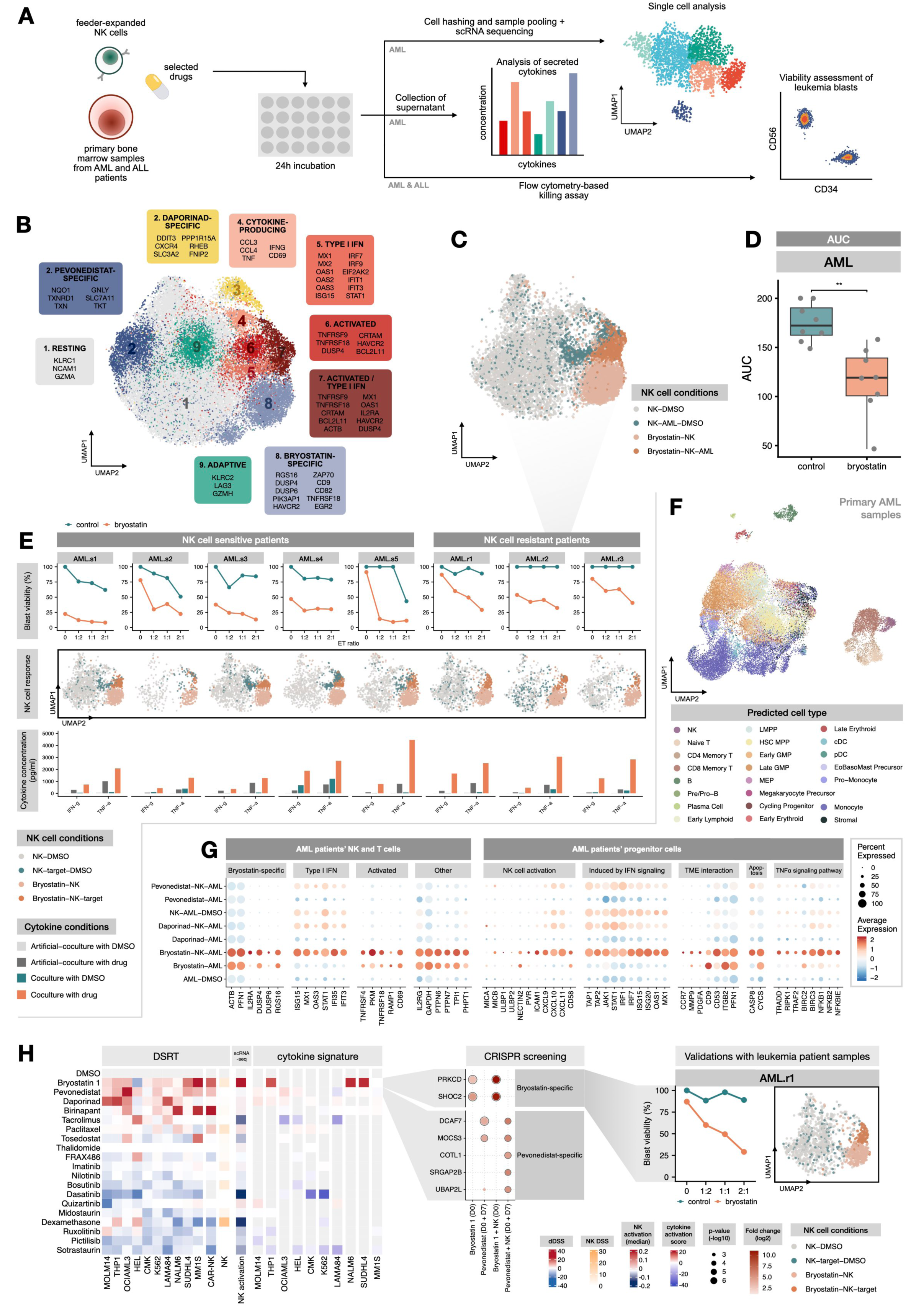
Validation experiments using patient bone marrow samples from AML patients and integrated data. (A) Workflow of validation experiments with primary AML and ALL bone marrow (BM) samples treated with expanded NK cells and either bryostatin 1 (10 nM) or pevonedistat (100 nM). (B) UMAP of expanded NK cells from AML patient sample validation experiments combined with NK cells from cell line experiments, including genes specific to each cluster. (C) UMAP of expanded NK cells from AML patient sample validations, indicating the distribution of cells within the UMAP depending on treatment condition. (D) Box plot of area-under-the-curve (AUC) values from AML patient sample validation experiments. (E) Plot with integrated data from AML patient sample validations, including flow-based killing assay readouts (top), scRNA seq of expanded NK cells cultured with AML patients’ BM samples (middle) and measurements of secreted cytokines (IFN-γ and TNF-α, bottom). (F) UMAP of AML patients’ cells, reflecting the predicted cell types. (G) Dot plot of selected genes found relevant to the function of NK cells in previous experiments, indicating their expression in AML patients’ NK and T cells (left). On the right, the expression of genes related to the interaction with NK cells in AML patients’ progenitor cells, shown as a dot plot. (H) Integrated plot, summarizing findings from the study. Heatmap (left) includes DSRT data from validated drugs in combination with NK cells and CAR NK cells, NK cell toxicity, NK cell activation scores from scRNA-seq, and cytokine signatures, which incorporates the log2 fold changes from selected cytokines (TNF-α, IFN-γ, MIP-1a, MIP-1b, GM-CSF, IP-10). Genome-scale CRISPR results (middle) summarize the most significant gene perturbations. Results from validations with leukemia patient samples highlight the impact of combining expanded NK cells with bryostatin 1 (right).

Using scRNA-seq, we identified drug-specific changes in the NK cell transcriptome with similar signatures observed in cell line experiments (Figure 7B, S7B and Table S4C, S4D). NK cells cultured with bryostatin 1 alone transitioned into the bryostatin-specific cluster, where genes specific to bryostatin 1 treatment (*RGS16, ACTB*, *PFN1*, *DUSP6*) were upregulated in addition to markers of activated NK cells, such as *HAVCR2* (TIM-3) (Figure 7B, 7C, S7B, S7C, Table S4C and S4D).

Of the eight primary bone marrow samples, five were sensitive to NK-cell-mediated killing, while three were resistant. Strikingly, results from killing assays showed that bryostatin 1 significantly improved the sensitivity of leukemic blasts to NK cells (p = 0.0078) even in patient samples resistant to NK cell treatment alone (Figure 7D). In NK cell-sensitive AML samples, in combination with bryostatin (10 nM), we achieved up to 90% killing of leukemic blasts at a 2:1 ET ratio and were able to reach 75% killing even in NK cell-resistant patients (Figure 7E, Table S7E).

In the case of ALL, both patients tested in our flow cytometry based killing assay were resistant to NK cells (Figure S7D). However, when combining NK cell treatment with bryostatin 1, we observed up to 50% killing off ALL blasts at 2:1 ET ratio (Figure S7D, Table S7F). Although statistically insignificant due to the low number of patients, the difference between controls and bryostatin 1-treated conditions was notable.

Integrating our findings from killing assays, transcriptomics and secretome analyses allowed us to identify differences in NK cell responses between NK cell-sensitive and resistant AML patients (7E). In NK cell-sensitive patients, NK cells co-cultured with primary AML cells were found mostly in the activated and type I interferon clusters. However, in NK cell-resistant patients, NK cells co-cultured with primary leukemia blasts remained primarily in the resting cluster. When adding bryostatin 1 to the co-culture, NK cells co-cultured with both NK cell-sensitive and resistant AML samples transitioned into the type I interferon and activated clusters, matching the strong responses observed in the killing assays. Moreover, cytokine measurements from matched experiments revealed a strong increase in IFN-γ and TNF-α secretion upon co-culture with bryostatin 1. The contrast was particularly striking in NK cell-resistant patients, where control co-cultures showed virtually no detectable IFN-γ or TNF-α, yet bryostatin 1 treatment elicited a robust cytokine response (Figure 7E).

In addition to effects observed in expanded NK cells upon combination with bryostatin 1, we also identified changes in AML patients’ lymphocytes and progenitor cells (Figure 7F, S7E and S7F). Interestingly, bryostatin 1 treatment caused transcriptional changes to patients’ NK and T cells similar to those observed in expanded NK cells, with further strengthened type I interferon and activated signals upon co-culture with expanded NK cells (Figure 7G and S7G). In AML patients’ progenitor cells upon treatment with expanded NK cells and bryostatin 1, we observed increased expression of markers involved in NK cell activation, interferon signaling, tumor microenvironment interaction, apoptosis, and TNF-α signaling (Figure 7G and S7H).

Altogether, these findings corroborated our cell line-based results and provided further insight into the mechanisms underlying the enhanced cytotoxic capacity of NK cells following bryostatin 1 treatment (Figure 7H). Importantly, this combination not only augmented NK cell cytotoxicity in primary leukemia samples but also mitigated resistance to NK cell-based therapies.

## DISCUSSION

In this study, we demonstrate how multimodal immunopharmacologic screening can systematically identify drug-mediated mechanisms that reprogram both immune effector function and tumor-cell susceptibility to enhance NK cell cytotoxicity. By profiling more than 500 compounds across leukemia, lymphoma, and myeloma cell lines, we identified bryostatin 1 and pevonedistat as the most potent enhancers of NK cell-mediated killing, each acting through distinct, multi-layered mechanisms. Through the integration of high-throughput functional drug screening, single-cell transcriptomics, cytokine profiling, and genome-wide CRISPR perturbation analysis, we uncovered changes in NK cell activation states, cytokine signaling pathways and tumor priming mechanisms that together explain the synergy observed between NK cells and drugs.

Beyond identifying individual compounds, our work establishes the high-throughput immune cell drug sensitivity and resistance testing platform as a powerful and scalable framework for discovering pharmacologic modulators of immune cell cytotoxicity. Combining functional killing assays with molecular profiling and genetic screening enables rapid prioritization of compounds that enhance or impair immune cell activity while revealing the mechanisms of responsiveness. This approach, using NK cells, builds on prior work applying similar discovery pipelines to CAR T cells^39^, highlighting the broader translational potential of multimodal immunopharmacologic screening across immune effector modalities.

A key finding of this study is the demonstration that effective NK cell potentiation arises from rewiring across both immune and tumor compartments, rather than isolated modulation of a single pathway in either cell type. In the case of bryostatin 1, we identified PKC signaling as a key regulator of NK cell activation. Single-cell transcriptomic analyses revealed transcriptional effects consistent with cytoskeletal remodeling, vesicular trafficking and immune synapse formation - key processes for efficient secretion of lytic granules and cytokines at the immunological synapse^18^. Importantly, cytokine profiling showed amplification of IFN-γ, TNF-α, and other cytokines’ secretion, indicating a shift towards a highly responsive and potentially self-reinforcing NK activation state. These effects occurred without upregulation of cytotoxic effector genes, suggesting that functional enhancement is driven mainly by increased secretory efficiency and signaling sensitivity rather than overexpression of effector genes.

Previous studies have also underscored the importance of PKC-θ in NK cell function, which is required for sustained signaling, transcriptional activation, and IFN-γ secretion^27,29^. Other PKC activators, such as ingenol 3,20-dibenzoate (IDB), enhance IFN-γ production and degranulation in NK cells exposed to non-small cell lung carcinoma cells^46^. Building on this knowledge, our results show that PKC activation not only increases IFN-γ secretion but also triggers transcriptional activation of multiple genes critical for NK cell anti-tumor activity, including upregulation of activating receptors and enhancement of cytoskeletal machinery, leading to more efficient effector molecule release.

Bryostatin 1 did not act solely on NK cells. Reflecting earlier findings from our group on cancer cells’ sensitivity and resistance mechanisms to NK cells^32^, tumor-cell transcriptomic profiling revealed increased expression of activating ligands, enhanced interferon signaling, and upregulation of genes supporting immune interaction and synapse formation, indicating a pre-sensitized tumor state that lowers the threshold for NK cell-mediated killing. Genome-wide CRISPR screening further identified PKC-δ as a genetic determinant of tumor sensitivity under bryostatin 1 treatment, suggesting PKC signaling has a role in both strengthening immune effector functions and increasing tumor cell vulnerability. Together, these findings support a two-sided model where PKC activation amplifies NK cell cytotoxic capabilities while priming tumor cells for improved responsiveness to immune-mediated killing.

The role of PKC in modulating NK cell-tumor interactions has been recognized in other settings, including increased immunogenicity of CLL cells^47–49^ and modulation of monocyte apoptosis via caspase-3 phosphorylation^50^. Our genome-wide CRISPR screening further highlighted PKC-δ, as an important determinant of sensitivity to NK cells upon bryostatin 1 treatment, suggesting that specific PKC isoforms may contribute both to NK cell activation and to enhanced target cell susceptibility. This genetic evidence complements our transcriptomic data, particularly as sequencing likely captured only those tumor cells surviving the potent combination of bryostatin 1 and NK cells.

While PKC-θ has been implicated in NK cell function previously, the mechanistic breadth revealed here, spanning enhanced receptor expression, cytoskeletal reprogramming, and tumor cell sensitization, extends our understanding of how PKC activation potentiates NK cell cytotoxicity. Importantly, this strategy increased NK cell-mediated killing of primary leukemic blasts, including cells from both NK-sensitive and NK-resistant patients, highlighting PKC activation as a promising avenue for therapeutic manipulation of NK cell function.

Although bryostatin 1 demonstrated potent activity, its broader clinical use may be limited by manufacturing complexity and scarce availability^51^. However, the development of bryostatin analogs^52^ offers a promising avenue to exploit PKC activation to improve NK cell-based immunotherapy. Beyond PKC signaling, our platform also identified NAE inhibition as another powerful route to enhance NK cell cytotoxicity, with pevonedistat emerging as a key modulator. In NK cells, single-cell transcriptomic profiling revealed increased expression of genes downstream of NRF2 activation, suggesting an improved capacity of NK cells to cope with oxidative stress, a known inhibitor of NK cytotoxicity in tumot contexts^33,53,54^. Notably, similar findings have been reported in ovarian cancer, where NRF2 activation with RTA-408 (omavexolone), an FDA-approved drug for Friedreich’s ataxia, restored cytotoxic function in primary NK cells^30^. Our results reflect these observations in the context of hematological malignancies and demonstrate that pevonedistat can drive a comparable NRF2-activated phenotype in NK cells. In addition, our screen identified daporinad as another metabolism-modulating compound that enhanced NK cell cytotoxicity, characterized by increased IFN-γ production and upregulation of glycolytic gene programs, underscoring the role of metabolic regulation in NK cell function.

In tumor cells, pevonedistat altered transcriptomic programs related to TNF signaling and apoptotic susceptibility. This aligns with previous reports showing that pevonedistat can synergize with TNF-α^55^. Because TNF-α is a key effector secreted by NK cells, this provides a plausible mechanism for the observed synergy, although the underlying processes have not been fully defined. Our analysis revealed that NAE inhibition shifted the balance of TNF receptor signaling in cancer cells, promoting pro-apoptotic pathways while suppressing NF-κB-dependent survival signaling. This is consistent with the established role of ubiquitination, disrupted by NAE inhibition, in regulating TNF signaling cascades^56^. Together, these findings suggest that pevonedistat sensitizes tumor cells to NK cell-derived TNF-α by interfering with ubiquitination-dependent checkpoints, thereby amplifying NK cell cytotoxicity as observed in our DSRT assays.

Overall, our results indicate that NK-drug synergy emerges from multi-axis rewiring of both NK and tumor cells, including enhanced immune activation, strengthened cytokine signaling, improved metabolic fitness of NK cells and increased apoptotic priming of cancer cells. Unlike therapeutic strategies that rely on a single molecular vulnerability, the NK-drug combinations identified here create partially independent pressures on tumor survival, potentially increasing the barrier for cancer cells to develop immune evasion mechanisms or therapeutic resistance. To evade NK cell and drug-mediated killing, tumor cells would need to simultaneously counteract increased immune effector functions and immune-sensitization mechanisms, creating a substantially more restricted adaptive landscape.

In conclusion, by integrating functional screening with multimodal molecular profiling, our high-throughput DSRT platform can systematically identify drug-mediated pathways that reprogram NK cell-tumor interactions, providing mechanistic insight and highlighting actionable targets. This supports rational combinatorial strategies that potentiate immune cell function while constraining tumor adaptation and resistance.

## METHODS

### Cell lines and expanded NK cells

All cancer cell lines were obtained from commercially available sources and low passage numbers were used to reduce the risk of subclonal selection. All cell lines were cultured in R10 medium, consisting of RPMI-1640 with 10% FBS, 2 mM L-glutamine, and 100 U/ml penicillin with 100 mg/ml streptomycin. All cultures were incubated at 37°C with 5% CO_2_.

Cells used for drug sensitivity and resistance testing were transduced to express luciferase. Luciferase-expressing MOLM-14, THP-1, OCI-AML-3, K562, CMK, LAMA-84, NALM-6 and MM1S were generated using pLenti PGK V5-Luc Neo (w623-2) construct as described previously^39^. HEL was transduced using EF1a-Luciferase (firefly)-2A-GFP (puro) lentivirus (Amsbio, LVP437) as described earlier^39^. Luciferace-expressing SU-DHL-4 was generated as described previously^39^. For scRNA sequencing experiments, parental cell lines were used.

All cell lines were STR profiled and tested for mycoplasma using the MycoAlert kit (Lonza).

### Primary NK cell expansion

NK cells were expanded from buffy coats of healthy donors, obtained from the Finnish Red Cross Blood Service, using K562-mbIL21-41BBL (CSTX002 Kiadis, Sanofi) feeder cells as described previously^57^. In short, peripheral blood mononuclear cells (PBMCs) were isolated from buffy coats using Ficoll-Paque gradient centrifugation. All material was handled in accordance with the Declaration of Helsinki and the Helsinki University Hospital ethical committee, statement number HUS/2215/2018. Expansions were started by suspending 5 million PBMCs, together with 10 million 100 Gy irradiated K562-mbIL21-41BBL (CSTX002 Kiadis, Sanofi) feeder cells, into 40 ml R10 supplemented with 10 ng/ml recombinant IL-2 (R&D Systems, 202-IL-050). Cells were passaged twice per week and feeder cells were added at a 1:1 ratio every 7 days. After 14 days of expansion, NK cells were purified using the NK Cell Isolation Kit (Miltenyi) and frozen in freezing media (90% FBS, 10% DMSO). NK cells were thawed 3 days prior to experiments and cultured in R10 supplemented with 10 ng/ml recombinant IL-2 (R&D Systems, 202-IL-050).

### CAR NK cell production

NK cells were preactivated with IL-15 (140IU/ml, Miltenyi) and IL-2 (500IU/ml, Miltenyi) for 3 days prior to transduction. NK cells were transduced with BaEVRLess pseudotyped lentiviral vectors (LVs) in NK cell expansion medium with a final cell density of 500,000 cells/ml using a multiplicity of infection (MOI) of 10. Vectofusin-1 (10 μg/ml, Miltenyi Biotec) was added simultaneously with the LVs after which NK cells were centrifuged at 400 g for 2 hours at room temperature. Following transduction, the LVs were washed away 24 hours after the transduction using PBS (day 4). Subsequently, the transduced cells were co-cultured with feeder cells (K562-mbIL-15-41BBL) at a 1:5 NK:feeder ratio from day 4 until day 12. On day 12, an additional dose of feeder cells (1:10 NK:feeder ratio) was added. After expansion, CAR-NK cell phenotype was analysed with flow cytometry (day 18) and cells were cryopreserved (day 19) in MACS ® Freezing Solution (Miltenyi). CAR NK cells were thawed 3 days prior to experiments and cultured in R10 supplemented with IL-15 (140IU/ml, Miltenyi) and IL-2 (500IU/ml, Miltenyi).

### Effector-target ratio optimization

Prior to performing high-throughput drug sensitivity and resistance testing screens on target cells with NK-drug combinations, an effector-target (ET) ratio at which 50% inhibition of target cell viability by NK cells alone had to be determined. This allowed us to determine both enhancing and inhibiting effects of drugs on NK cells.

Expanded NK cells were thawed 3 days prior to the experiment and cultured in media containing IL-2 (10 μg/ml) at 1.0 x 10^6^ cells/ml. Target cells were thawed 1 day prior to the experiment and seeded at appropriate concentrations according to supplier’s instructions. On the day of the experiment, media was changed from the NK cell suspension and replenished with fresh R10 and the concentration was adjusted to 2.0 x 10^6^ cells/ml. The NK cell suspension was serially diluted to reach concentrations of 1.0 x 10^6^, 0.5 x 10^6^ and 0.25 x 10^6^ cells/ml. Media from target cells was also changed, and cells were resuspended in R10 at 1.33 x 10^6^ cells/ml. In 384-well plates, 15 μl of target cell suspension was then plated, followed by 10 μl of either R10 or appropriate NK cell suspension to obtain ET ratios of 0:1, 1:1, 1:2, 1:4 and 1:8. For each ET ratio and target cell line, 6 replicates were plated for statistically significant results. Plates were incubated for 24 h at 37°C, after which luciferase luminescence was detected following the addition of 25 μl/well of ONE-Glo (Promega) and readout using PHERAstar FCX microplate reader (BMG Labtech). Viabilities were calculated based on luminescence readouts using R.

### Drug sensitivity and resistance testing (DSRT)

Drug sensitivity and resistance testing (DSRT) of 527 approved and investigational oncology compounds at five different concentrations (FO5A FIMM oncology library) was used for high-throughput evaluation of their effects on NK cell cytotoxicity and their impact on NK cell viability (Table S1A).

Luciferase-expressing target cells were cultured in R10 and the cell suspension was adjusted to 2.0 x 10^6^ cells/ml. NK cells were thawed 3 days prior to the experiment and cultured in media containing IL-2 (10 μg/ml) at 1.0 x 10^6^ cells/ml. On the day of the experiment, the media was changed to R10 and the cell concentration was adjusted to reach the appropriate ET ratio used with each cell line. Drug plates were prepared by plating 5 μl of R10 using a multi-mode dispenser (BioTek, Multiflo FX with stacker, 5 µL RADTM cassette) to each well to dissolve the pre-plated drugs, followed by short centrifugation and 10 min shaking at 450 rpm, variable speed. Target cells were then plated, 10 μl/well, followed by 10 μl/well of NK cell suspension or R10, depending on the plate set. For each cell line, both an NK cell set and an R10 set were plated, to obtain data on the effects of the drug alone and in combination with NK cells. Finally, after plating cell suspensions, plates were put on a shaker for 5 mins, 450 rpm, variable speed. Plates were then incubated for 24 h at 37°C, after which luciferase luminescence was detected following the addition of 25 μl/well of ONE-Glo (Promega) and readout using PHERAstar FCX microplate reader (BMG Labtech).

The same platform was used to assess the effects of drugs on NK cell viability by adding 15 μl of R10 and 10 μl of NK cell suspension (at 2.0 x 10^6^ cells/ml) per well. Following 24 h incubation, the viability of NK cells was assessed using CellTitre-Glo (Promega) and readout using PHERAstar FCX microplate reader (BMG Labtech).

### Drug screening data analysis

Luminescence readouts from each well were processed to generate summarized measurements of all compounds in the library under each experimental condition using a pipeline based on the Breeze web software developed at the Institute of Molecular Medicine in Finland (FIMM)^58^. We adapted the pipeline to our in-house plate layout and co-culture assay format. Breeze generates quality control metrics and visualizations for each plate, which were used to assess data quality. In addition, Breeze integrates dose-response profiles to quantify compound activity using the drug sensitivity score (DSS)^59^. The DSS is calculated based on the area under the dose-response curve (AUC), providing a standardized and more comparable measure of drug response rather than single-point parameters such as the IC₅₀ or EC₅₀^60^.

When both control and test cell populations are available, Breeze calculates differential drug sensitivity score (dDSS) results, quantifying selective responses in test cells relative to positive and negative controls. In this study, our control data consisted of cancer cells cultured alone, while test conditions contained cancer cells co-cultured with NK cells. The dDSS was calculated as the difference between the co-culture DSS and the average DSS of cancer cells having been treated with the drug alone. Compounds showing activity only at single concentrations or displaying unreliable dose-response curves were excluded from the analysis. Drug classes with multiple active compounds impacting cytotoxicity were highlighted as being of particular interest.

### Validation drug screening

For validating the results from high-throughput DSRT, a custom panel consisting of 36 compounds at 9 concentrations was designed (Table S2A). Compounds were selected by first excluding those with high toxicity to NK cells (DSS > 8), followed by selecting compounds with a cumulative absolute dDSS ≥ 60 across all 10 cell lines, or an absolute dDSS ≥ 20 in at least one cell line. This approach enabled the identification of both strong modulators of NK cell cytotoxicity and compounds with more moderate but consistent effects across multiple cell lines and disease types.

Validation experiments were carried out with NK cells from 3 different healthy donors representing different HLA-types, expanded as described earlier. In addition, CD19-targeting CAR NK cells generated from one healthy donor were tested against a CD19-positive cell line (NALM-6).

NK cells were thawed 3 days prior to the experiment and cultured in media containing IL-2 (10 μg/ml) at 1.0 x 10^6^ cells/ml. On the day of the experiment, the media was changed and replaced with R10 to reach an appropriate cell suspension concentration based on the ET ratio used for each cell line. Target cell suspensions were adjusted to 2.0 x 10^6^ cells/ml density in R10. Pre-plated drugs in 384-well plates were first dissolved by plating 5 μl of R10 using a multi-mode dispenser (BioTek, Multiflo FX with stacker, 5 µL RADTM cassette), followed by short centrifugation and 10 min shaking at 450 rpm, variable speed. 10 μl of target cell suspension was then plated into each well, followed by 10 μl of NK cell suspension or R10. Finally, after plating cell suspensions, plates were put on a shaker for 5 mins, 450 rpm, variable speed. Plates were then incubated for 24 h at 37°C, after which luciferase luminescence was detected following the addition of 25 μl/well of ONE-Glo (Promega) and readout using PHERAstar FCX microplate reader (BMG Labtech).

### Multiplexed single-cell RNA sequencing

#### Experiments and multiplexing

Single-cell RNA sequencing was used to investigate the effects of selected drugs on NK cells and target cells, both individually and in co-culture, to gain deeper insight into their impact on cellular phenotypes and underlying biological mechanisms. Drugs of interest were selected for further analysis based on a combination of scientific relevance and performance in our high-throughput DSRT screen and validation assays (Table S3A). Specifically, compounds showing favorable profiles in differential drug sensitivity and low toxicity to NK cells were prioritized. Drug concentrations were selected based on DSRT results, specifically choosing the dose that showed the greatest difference between drug-only and drug+NK conditions without causing complete inhibition of target cells.

Single-cell RNA sequencing was performed using unique oligonucleotide barcoded antibodies, which allowed us to pool cells from multiple conditions into each sample sent for sequencing while being able to identify which cell originated from which condition in the experiment (Table S3D).

Parental cancer cell lines and expanded NK cells were thawed 3 days prior to the experiment and cultured as previously described for the DSRT experiment. NK cells were expanded from a single healthy donor, and all cells used in the experiments originated from the same expansion batch. For each experiment, 2 experimental batches were sent for sequencing at a time, consisting of 12 conditions each. 250,000 target cells were plated on 24-well plates either alone or with NK cells at appropriate ET ratios based on optimization experiments, and a matching number of NK cells were plated alone in conditions where the effect of drugs on NK cells were evaluated alone. To drug conditions, 5 μl of drug suspension at appropriate concentration was added, in addition to R10, to make up to a total volume of 1 ml per well. For control conditions, cells were cultured with DMSO concentrations matching those of drug conditions in which DMSO was used as solvent. Plates were then incubated for 24 h at 37°C.

Following 24 h incubation, samples were prepared for sequencing. Cells were transferred to 15 ml Falcon tubes and kept on ice. Samples were then centrifuged for 5 mins at 300 rpm, and 2 x 100 μl samples of the supernatant were collected and stored at -80°C for later use in cytokine analyses. Cells were then washed 3 times with 10 ml of cold PBS and centrifuged at 300 rpm for 5 minutes in between. After the final wash with PBS, the supernatant was carefully removed and cells were resuspended in 100 μl of cold staining buffer (2% BSA / 0.01% Tween in PBS) and transferred to FACS tubes placed in ice. 10 μl of blocking reagent (Human TruStain FcX, Biolegend) was then added to each sample, tubes were vortexed and incubated at 4°C for 10 mins. Following incubation, 1 μg (2 μl) of unique cell hashing antibody (TotalSeq A, Biolegend) was added to each sample and a record was made of which antibody was used for which sample (Table S3D). Tubes were vortexed and incubated at 4°C for 30 mins. Cells were then washed with 3 ml of cold staining buffer 5 times, with centrifugation of 300 rpm at 4°C for 5 min in between washes. After the final wash, each sample was resuspended in 100 μl staining buffer and samples were merged. The merged samples were centrifuged at 300 rpm, 4°C, for 5 mins and the supernatant was removed. Cells were then resuspended in cold PBS containing 0.04% BSA and cell density was adjusted to 1500 cells/μl. Samples were kept on ice and taken for further processing.

#### scRNA-seq library preparation and sequencing

The Chromium Single Cell 3’ RNASeq run and library preparations were done using the 10X Genomics Chromium Next GEM Single Cell 3’ Gene Expression v3.1 Dual Index chemistry, following the manufacturer’s instructions with modifications described earlier^32,61^. Targeting approximately 1,100-1,200 cells per condition for sequencing, approximately 15,000 cells from samples containing a pool of 13 hashed samples from different conditions were loaded into each 10X GEM well. Libraries were sequenced on an NovaSeq 6000 (Illumina) system using the following read lengths: 28 bp (Read 1), 10 bp (i7 Index), 10 bp (i5 Index), and 90 bp (Read 2).

Sequencing data were processed using the 10x Genomics Cell Ranger software (v5.0.1). FASTQ files were generated using the cellranger mkfastq pipeline with Illumina bcl2fastq v2.2.0, and gene expression matrices were obtained using the cellranger count pipeline with alignment to the GRCh38 human reference genome. Data from multiple samples were combined using the cellranger aggr pipeline to generate a unified gene-barcode matrix.

Cell hashing was performed according to the manufacturer’s instructions (BioLegend protocol for TotalSeq-A antibodies and cell hashing with 10x Single Cell 3’ Reagent Kit v3.1). Hashtag oligonucleotide (HTO) count matrices were generated using the CITE-seq-Count tool^61^.

### Single-cell RNA sequencing analysis

#### Quality control and pre-processing

The R package Seurat (≥v4.3.0)^62^ was used for further scRNA-seq data processing. Cell types were annotated using SingleR to distinguish NK and target cells. We filtered out all cells with mitochondrial gene counts, number of detected genes, or UMI counts outside the ranges defined in Table S3F. Hashtag oligonucleotide (HTO) demultiplexing was performed on centered log-ratio-normalized UMI counts to assign cells to samples, separately for the two plates. This was done with an adapted version of Seurat’s HTODemux function followed by an outlier removal, which is discussed in the next section. Only cells classified as singlets were considered for further analyses. A few cells considered doublets, as they expressed both genes associated with NK and target cells, were manually removed. The transcriptome data was log-normalized, and the highly variable genes were calculated with the FindVariableFeatures function using the vst selection method in Seurat.

#### Adapted HTODemux and outlier removal

HTODemux performs initial clustering with k-medoids, and for each HTO uses the cluster with lowest expression to estimate the background expression. Here, the two clusters with lowest expression were used instead to get a better estimate of the background expression especially in cases where the first cluster consists of only a few cells. In addition, we set two different parameter values for the positive quantile value (pq), one for detecting multiplets, and another for detecting negatives from the remaining cells (Table S3F). This allowed us to better control the number of cells considered as multiplets and negatives. The number of expected doublets was computed based on the Chromium Single Cell 3’ Reagent Kits User Guide (v3.1 Chemistry Dual Index) and the number of detected cells from each plate. When selecting the pq values, the number of predicted multiplets was compared to the number of expected multiplets to ensure the number of predicted multiplets was reasonable. From the cells classified as singlets, we further identified outliers using DBSCAN^63^. DBSCAN was applied separately to cells assigned to each HTO to identify the main cluster(s) first based on t-SNE^64^, and then on UMAP^65^ dimensional reduction spaces, computed with all 12 PCs. Cells outside these clusters were considered as outliers and removed.

#### Batch correction

To correct for batch effects across different experimental plates, we used a probabilistic framework using the deep generative modeling tool single-cell variational inference (scVI) (v 1.0.0)^66^. 2,000 highly variable genes (HVGs) identified by Scanpy using seurat_v3 flavour were used and cell cycle phase was corrected based on G2M and S scores calculated with Seurat’s CellCycleScoring function. With NK cells 50 scVI latent dimensions were used and for other data 30 latent dimensions were used. For analyses that focus on certain subsets, such as all NK cells or AML cell lines, batch correction was done separately for the selected cell subset in the same manner.

#### Visualization, clustering, cell type annotation and differential gene expression

The UMAP dimensionality reduction with default parameters was computed using RunUMAP and clustering was done using Seurat’s FindClusters and FindNeighbors functions, for both all computed scVi latent components were used. For expanded NK cells from cell line experiments clusters were identified with resolution 0.4 which resulted in 11 clusters, out of which three were combined to form the Resting cluster. Clusters for expanded NK cells from experiments with primary AML cells and NK cells from cell line experiments with the same drugs (bryostatin 1, daporinad, pevonedistat) and matching DMSO conditions were identified with resolution 1.3 which resulted in 18 clusters, out of which one low quality cluster was removed, two clusters were combined as Bryostatin-specific, and eight were combined to form the Resting cluster. The broad cell types of AML patient cells were annotated with BoneMarrowMap R package^67^. Differential gene expression (DEG) was computed with Seurat’s functions FindMarkers and FindAllMarkers using Wilcoxon Rank Sum test and adjusted p-values were bonferroni corrected using all genes in the dataset being evaluated. For DEG analysis of AML patient cells Naive T, CD4 Memory T, CD8 Memory T, and NK were considered as NK and T cells and HSC MPP, Early GMP, Late GMP, MEP, Megakaryocyte Precursor, Cycling Progenitor, EoBasoMast Precursor, and LMPP were considered as progenitor cells.

### Cytokine measurements and analysis

#### Sample collection and determining cytokine concentration

Supernatant samples were collected from the same samples later processed for single-cell RNA sequencing, except in the case of NALM-6 and SU-DHL-4, where drugs were selected based on top drug candidates across multiple disease types (Table S3B). From these samples, cytokine concentrations were measured using the Bio-Plex Pro™ Human Cytokine 27-Plex Assay (#M500KCAF0Y, BioRad) according to the manufacturer’s protocol. Samples were analyzed using the Bio-Plex 200 system (BioRad).

#### Analysis

Cytokine data analysis was performed using R. Background cytokine levels were subtracted from experimental samples using control measurements from tissue culture media containing either DMSO or the tested drugs.

To increase statistical power, NK cells treated with the same drug dose across different experiments were pooled and compared to corresponding control conditions. Statistical significance was assessed using the Wilcoxon signed-rank test. Fold changes were calculated by comparing cytokine levels in the supernatant of drug-treated NK cells to the average cytokine levels of DMSO-treated controls.

To assess the effect of co-culture, artificial co-culture cytokine values were calculated by summing cytokine concentrations from NK-only and target-only conditions for each treatment. These artificial values were then compared to true co-culture measurements to evaluate changes in soluble factor secretion upon drug treatment.

### Degranulation assay

#### Degranulation experiment

We established a flow cytometry-based degranulation assay to evaluate the effect of drugs on NK cell degranulation and activation (Table S7A). Experiments were started by thawing expanded NK cells three days prior to the experiment and culturing them as previously described. Target cell lines (MOLM-14, THP-1, and OCI-AML-3) were similarly thawed three days in advance and cultured in R10.

To a 96-well plate, NK-only, target-only, and co-culture conditions were plated, with and without drugs, also including wells dedicated for unstained samples. Into each well containing target cells, 50,000 cells/well were added. Based on earlier ET ratio optimizations, 50,000 target cells were added into NK-only and THP-1 co-culture wells (1:1 ET ratio), and 25.000 target cells were added into MOLM-14 and OCI-AML-3-containing wells (1:2 ET ratio). Appropriate amounts of drugs suspended into R10 were added to drug-containing wells (10 nM bryostatin 1, 100 nM pevonedistat). Prior to incubation, CD107a antibody was added into stained wells. The plate was shaken for 5 min at 450 RPM (variable speed), and placed in the incubator. After 1 h of incubation, GolgiStop was added and the plate was placed on a shaker for 5 min at 450 RPM (variable speed). Incubation was then continued for another 5 h.

#### Staining and flow cytometry

Following a total of 6h incubation, cells were washed with the staining buffer (PBS with 2 mM EDTA and 0.05% BSA) and stained for viability (Table S7A), followed by a 30 min incubation at 4°C in the dark. Samples were then washed with the staining buffer, followed by the addition of surface marker antibodies (Table S7A). Samples were then incubated for 15 min at room temperature in dark, followed by a wash with the staining buffer. Samples were then fixed using the Fixation/Permeabilization Kit (BD Cytofix/Cytoperm, 554714) and incubated for 20 min at 4°C in the dark. Samples were then washed twice using the staining buffer and resuspended into 50 μl staining buffer. The plate was covered from light and placed in 4°C overnight. On the following day, samples were washed and resuspended into fixation/permeabilization wash buffer, following the addition of intracellular antibodies (Table S7A). Samples were allowed to incubate for 30 min at room temperature in the dark. Samples were then washed with fixation/permeabilization wash buffer, and resuspended into 50 μl staining buffer. Samples were acquired using Novocyte Quanteon flow cytometer.

#### Analysis

Degranulation assay data were analyzed using FlowJo software (Becton Dickinson & Company (BD)) and further analysis was done using R.

### TNF-α blocking experiment with infliximab

To confirm the importance of TNF-α in the enhanced cytotoxic effect seen when combining pevonedistat with NK cells, we designed an assay making use of TNF-α blocking using the monoclonal antibody infliximab.

Expanded NK cells were thawed three days prior to the experiment and cultured as previously described. Target cell lines (MOLM-14 and OCI-AML-3) were similarly thawed three days in advance to ensure adequate recovery after thawing.

For the assay setup, 10 μl of target cell suspension (2.0 × 10⁶ cells/ml in R10 medium) was added to each well of a 384-well plate. For target cell-only controls, an additional 5 μl of R10 medium was added. In co-culture conditions, 5 μl of NK cell suspension was added at 1:2 effector-target (ET) ratio.

To each well, 5 μl of either R10 medium (vehicle control) or infliximab diluted in R10 was added, depending on the condition, resulting in a final infliximab concentration of 10 μg/ml. For drug treatment conditions, an additional 5 μl of drug solution was added to reach a 100 nM pevonedistat concentration. In control wells without drug treatment, 5 μl of R10 medium was added instead. All conditions were plated in triplicate.

### High-throughput flow cytometry and scRNA-sequencing validation experiments with patient samples

Main drug candidates were validated using patient samples from the iCAN cohort from which 8 AML patients’ and 2 ALL patients’ samples were used for experiments (Table S3C). ALL patients were selected from B-cell ALL cases with CD19-positivity as a criteria allowing for the use of our B-ALL panel in flow cytometry.

#### Co-culture experiment with primary leukemia samples and NK cells

Expanded NK cells from the same healthy donor used in cell line single-cell RNA sequencing experiments were thawed 3 days prior to the experiment and cultured as described earlier. Bone marrow (BM) samples from AML and ALL patients were thawed one day prior to the experiment. After thawing, BM samples were washed once with R10 medium and resuspended into 1 ml R10 medium. 50 μl of DNase (RNase-free DNase RQ1, Promega) was added to the cell suspension, following a 20 min incubation at 37°C. Samples were then passed through a 30 μm pre-separation filter and cells were washed with 10 ml of R10 medium. DNase-containing medium was then removed and cells were resuspended into R10 medium at 1.0 x 10^6^ cells/ml and cultured until the next day to allow for recovery post-thawing.

On the day of the experiment, the medium of both NK cell and patients’ cell suspensions were changed into fresh R10 medium. For samples to be sent for sequencing, 125,000 patients’ leukemic cells were plated with or without NK cells at a 1:2 ET ratio, both with and without drugs and DMSO as vehicle control. Drugs and their respective concentrations used are listed in Supplemental Table S3C. Also NK cells alone with and without drugs were plated. After plating into 96-well plates, plates were placed on a shaker for 5 mins at 450 rpm, variable speed, following 24h incubation 37°C.

In parallel with the sequencing experiment, a flow cytometry-based killing assay was performed to evaluate the effect of NK cell-drug combinations on patient leukemic blast viability. Patient cells (25,000 per well) were plated in a 96-well plate. For wells without NK cells, R10 medium was added, whereas NK cell-containing wells received the appropriate number of NK cells to achieve three effector-target (ET) ratios: 2:1, 1:1, and 1:2. Finally, either DMSO (vehicle control) or drugs at the concentrations specified in Table S3C. All conditions were prepared in triplicate. Plates were shaken for 5 min at 450 rpm (variable speed) and then incubated for 24 h at 37 °C.

#### scRNA-seq and flow cytometry

After the 24 h incubation, samples were prepared for multiplexed single-cell RNA sequencing as described earlier, and supernatants were collected for cytokine analysis (Table S3E). Samples from the killing assay were processed for flow cytometry. Cells were transferred to a 96-well v-bottom plate and centrifuged at 500 rpm for 5 min, after which the supernatant was removed. For surface staining, 25 μl of antibody master mix (2,310 μl staining buffer with antibodies listed in Tables S7B and S7C) was added to each well. Cells were resuspended and mixed by pipetting 10 times, then incubated for 20 min at room temperature protected from light. Plates were centrifuged for 5 min at 500 rpm, the supernatant was removed, and cells were washed with 100 μl PBS containing 0.1% BSA, followed by centrifugation under the same conditions. For viability staining, 25 μl of Annexin V/DRAQ7 master mix (2,415.4 μl distilled water, 275 μl Annexin V binding buffer, 4.6 μl DRAQ7, and 55 μl Annexin V) was added to each well, and cells were resuspended by pipetting 10 times. The plate was incubated for 10 min at room temperature, protected from light. Flow cytometry was performed on an iQue 3 instrument (Sartorius).

### CRISPR screening

#### Generation of Cas9-EGFP expressing cell line

The THP-1 cell line was transduced to for Cas9-EGFP using lentiCas9-EFGP plasmid (#63592, Addgene). First, 293FT cells (ThermoFisher Scientific, #R700-07) were suspended into 10 ml D10 media (DMEM, #E12-614F, Lonza; 10% FBS; 2 mM L-glutamine) at 3.8x10^5^ cells/ml. The lentivirus was produced by co-transfecting 293FT cells with 10 μg of lentiviral plasmid (#63592, Addgene), 7.5 μg psPAX2 lentiviral packaging plasmid (#12260, Addgene) and 2.5 μg pCMV-VSV-G lentiviral envelope plasmid (#8454, Addgene), in combination with 40 μl Lipefectamine 2000 (#11668019, Thermo Fisher Scientific) according to manufacturer’s instructions. 293FT cells were cultured at 70-80% confluency in D10 for 6 h of transfection, after which D10 was replaced with 15 ml bovine serum albumin (BSA) -containing D10 (D10 + 1% BSA). Lentiviral particles were harvested after 48 h, by collecting supernatant filtered through a 45 μm filter. Aliquots of supernatant were stored at -70°C.

Transduction of THP-1 cells was done by suspending 1.5 x 10^6^ THP-1 cells into 500 μl viral supernatant and 8 μg/ml Polybrene (#H9268-10G, Sigma-Aldrich) in a 24-well plate. The cell suspension was centrifuged at room temperature at 800 g for 2 h, following removal of viral supernatant. Cells were then cultured and passaged until a negative replication competent virus (RCV) test result could be obtained. Following a negative RCV test, cells were collected and the EGFP-positive cell population (THP-1) was sorted using a cell sorter SH800 (Sony) with the FITC channel. A THP-1 clone with high EGFP expression was selected for screening. Cas9 activity was confirmed by knock-out of beta-2-microglobulin (*B2M*), using sgB2M (5’-caccGCTACTCTCTCTTTCTGGCC-3’). Following knock-out, cells were stained with anti-HLAabc antibody (#561349, BD Biosciences) and analyzed with Sony SH800 cell sorter.

#### Library production

The amplified Brunello library (#73178, Addgene) in LentiCas9-Blast vector (#52962, Addgene) was a gift from E. Akimov (University of Helsinki) and was produced as previously described (Awad et al. 2024). To generate the lentiviral library, 8.9 x 10^6^ 293FT cells/175 cm^2^ flask were seeded the day before transfection in 23.3 ml of DMEM supplemented with 10 % FBS and 2mM L-glutamine. An hour before transfection the medium was replaced with 10.1 ml of Opti-MEM (#31985062, Thermo Fisher Scientific). The cells were transfected with 15.6 mg of the plasmid library, 7.8 mg of pCMV-VSV-G, 11.7 mg of psPAX2, 155.6 ml of Plus Reagent (#11514015, Thermo Fisher Scientific) and 77.8 ml of Lipofectamine 2000 per flask according to manufacturer’s protocol. After six hours of incubation the medium was changed to 23.3 ml of DMEM supplemented with 1% BSA, 10 % FBS and 2mM L-glutamine. After 60 hours of incubation the cell debris was removed from the virus media by centrifugation and the supernatant was filtered with a 0.45 mm filter. Aliquots of the library were stored at -70°C.

#### Lentiviral transductions with sgRNA library

The human CRISPR KO pooled Brunello library^68^ lentivirus was produced as described earlier using the amplified Brunello plasmid library (#73178, Addgene). The optimal lentivirus concentration for transduction was determined by testing a range of virus volumes, ranging from 0-1000 μl of the Brunello virus and 8 μg/ml Polybrene, on 3 million THP-1-Cas9-EGFP cells in a 24-well plate. The plate was centrifuged for 2 h at 800 g, following removal of viral supernatant. Cells were then cultured with or without 1 μg/ml pyromycin (A11138-03, Thermo Fischer Scientific) for 8 days, starting at 48 hours post-transduction. Transduction efficiency was evaluated 144 h post pyromycin treatment using CellTitre-Glo (#G9243, Promega), with readouts obtained using FLUOStar (BMG Labtech) microplate reader. Virus volume of 75 μl, yielding a transduction efficiency of 18.8%, was then selected for large-scale transduction.

Based on optimized transduction conditions, 163 x 10^6^ THP-1-Cas9-EGFP cells were transduced as described earlier, using 4075 μl of Brunello library-containing viral supernatant. Cells were then antibiotically selected and cultured until a negative RCV testing could be obtained, following freezing of cells for later experiments.

#### Optimization of screening conditions

To find drug concentrations that would keep cell numbers near constant throughout the CRISPR screen, ranges of drug concentrations and amounts of NK cells with THP-1-Cas9-EGFP were tested. The tested conditions were selected based on the drug screen reported here, and we tested ET ratios of 0:1, 1:8, 1:4, 1:2, and 1:1, alongside pevonedistat concentrations of 1-50 nM and bryostatin 1 concentrations of 3-50 nM, and selected conditions that maximized differential survival while maintaining sufficient library representation. Cells were seeded at 0.5 x 10^6^ cells/ml in 5 ml of cell culture medium in 6-well plate wells and the drugs in dimethylsulfoxide (DMSO; #D2650, Merck) were added at appropriate concentrations with and without NK cells at different ET ratios. 0.2 % DMSO was used as a negative control. Drugs and NK cells were added at days 0 and 7. Viable cells were counted and passaged at 0.5 x 10^6^ cells/ml and, following the evaluation of THP-1 viability using flow-cytometry to identify GFP signal intensity. Based on results from the optimization experiments, the 1:2 ET ratio was selected, along with 3 nM bryostatin 1 and 50 nM pevonedistat.

#### CRISPR screen

Two replicate CRISPR screens with THP-1 were performed, two consisting of 50 nM pevonedistat treated conditions and one replica including 3 nM bryostatin 1. Prior to the screen, THP-1 cells transduced with the Brunello library were thawed and expanded to reach a cell count necessary to perform the screen. The drugs and the 0.2 % DMSO control were added at optimized concentrations to 50 x 10^6^ cells in 100 ml in 175 cm^2^ cell culture flasks, with 25 x 10^6^ NK cells added to NK cell-containing conditions. Maximum of 50 x 10^6^ cells of each treatment were passaged every two or three days at 0.5 x 10^6^ cells/ml. Drug or DMSO and/or NK cells were added at days 0 and 7 of the screen. In the case of bryostatin 1, the drug and NK cells caused significant killing of THP-1 cells and hence only 1 dose of both NK cells and bryostatin 1 were added at day 0. Cell cultures were maintained for 14 days in total after which cell pellets containing a maximum of 100 million cells were harvested. The pellets were stored at -20°C.

#### Next generation sequencing

DNA was extracted from frozen pellets containing 3.5 x 10^7^ using Blood & Cell Culture Maxi Kits (Qiagen) and processed according to manufacturer’s instructions. DNA concentration was quantified using fluorometric quantification (Qubit, Thermo Fischer Scientific). Samples then went through a two-step PCR protocol as previously described^32,69^. Amplified samples were sequenced with Illumina NovaSeq 6000 (Illumina).

#### Analysis

CRISPR data were analyzed using MAGeCK v0.5.9.5^70^. Reads from the forward (R1) direction were aligned with the Brunello library sgRNA sequences using mageck count function with default parameters. To compare conditions, MAGeCK test analysis was performed on the sgRNA count matrices using the mageck test function with default parameters. For comparisons across conditions in pevonedistat treated replicas, the mageck test function with paired function was used. Based on gene summary files generated with MAGeCK, further analyses were done using R.

#### Data availability

All processed and raw sequencing data will be made publicly available upon publication of this work.

## ACKNOWLEDGEMENTS

We thank Drs. Karl-Johan Malmberg, Päivi Ojala, Shady Awad, Jani Huuhtanen, and Francesca De Lorenzo for insightful comments and support with our work. We also thank Sanna Timonen for help with CRISPR screen design and analysis. The authors would also like to thank healthy donors and patients for their generous contributions. The authors acknowledge the help and services received from the The Helsinki Institute of Life Sciences (HiLIFE), FIMM High-throughput Biomedicines, Single Cell Analytics, and Genomics Sequencing units; the Biomedicum Flow Cytometry Core; the Finnish Hematology Registry and Clinical Biobank (FHRB), and the Finnish Red Cross Blood Service. Authors also acknowledge the computing resources provided by CSC (IT Service for Science Ltd.). Portions of the manuscript were edited for clarity and style using an AI-based language model (ChatGPT 5.0, OpenAI). The study was supported by grants received from the Jane and Aatos Erkko Foundation, Leukemia and Lymphoma Society, Cancer Foundation Finland, Academy of Finland, Sigrid Juselius Foundation, the Gyllenberg Foundation, State funding for University-level Health Research in Finland, and HiLIFE fellow funds. J.B. was supported by grants from Emil Aaltonen Foundation, Tor, Joe & Pentti Borg Foundation, Ida Montin Foundation, The Paulo Foundation, The Finnish Medical Foundation, The Finnish Hematology Association, The Blood Diseases Research Foundation and Juhani Aho Foundation for Medical Research. T.A. was supported by iCAN Digital Precision Cancer Medicine Flagship (iCAN-MULTIDRUG). CSM and SM are supported by Blood Cancer United (formerly The Leukemia & Lymphoma Society) and by a grant from the NIH (R01 CA276156). CSM is also supported by the Ludwig Center at Harvard, Myeloma Solutions Fund, de Gunzburg Myeloma Research Fund, Cobb Family Myeloma Research Fund, and the Oliver S. and Jennie R. Donaldson Charitable Trust.

## AUTHORSHIP CONTRIBUTIONS

J.B. designed and conducted experiments, analyzed and interpreted results, and wrote the manuscript; E. Jokinen processed, analyzed and helped interpret scRNA-seq data; P.N. helped design experiments, and conducted drug screening and scRNA-seq experiments; D.D. and A.I. analyzed drug profiling data; J.K. and H.L. conducted drug screening experiments, helped with scRNA-seq experiments, NK cell expansions, and prepared luciferase-expressing cell lines; S.D. helped with CRISPR screens; H.D. provided assistance with leukemia patient sample experiments and NK cell expansions; E. Järvelä generated CD19 CAR NK cells; E.S. and A.N. performed scRNA-seq library preparations; K.M. and T.H. assisted with CRISPR screens; L.T. provided assistance with drug profiling; M.M., D.S., and S.Y.H. provided advisory soles; E.L. provided assistance with scRNA-seq analysis; D.L. provided assistance with NK cell expansion methods; M.K. and H.G. supervised work related to CAR NK cells experiments; T.A. designed and supervised drug profiling data analyses; M.H. performed cytokine measurements; C.M. provided advisory roles and helped interpret results; S.G. helped design and perform CRISPR screens, helped with NK cell expansions, and helped interpret results; O.D. and S.M. supervised the project, led the design, analysis and interpretation of data, and contributed to the writing of the manuscript.

## DECLARATION OF INTERESTS

TA has received unrelated research funding from Mobius Biotechnology GmbH. C.S.M. has served on the Scientific Advisory Board of Adicet Bio and discloses consultant/honoraria from Genentech, Nerviano, Secura Bio and Oncopeptides, and research funding from EMD Serono, Karyopharm, Sanofi, Nurix, BMS, H3 Biomedicine/Eisai, Springworks, Abcuro, Novartis and OPNA. SM has received unrelated research funding from Novartis, Bristol Myers Squibb and Pfizer.

---

**Supplemental Figure 1:**
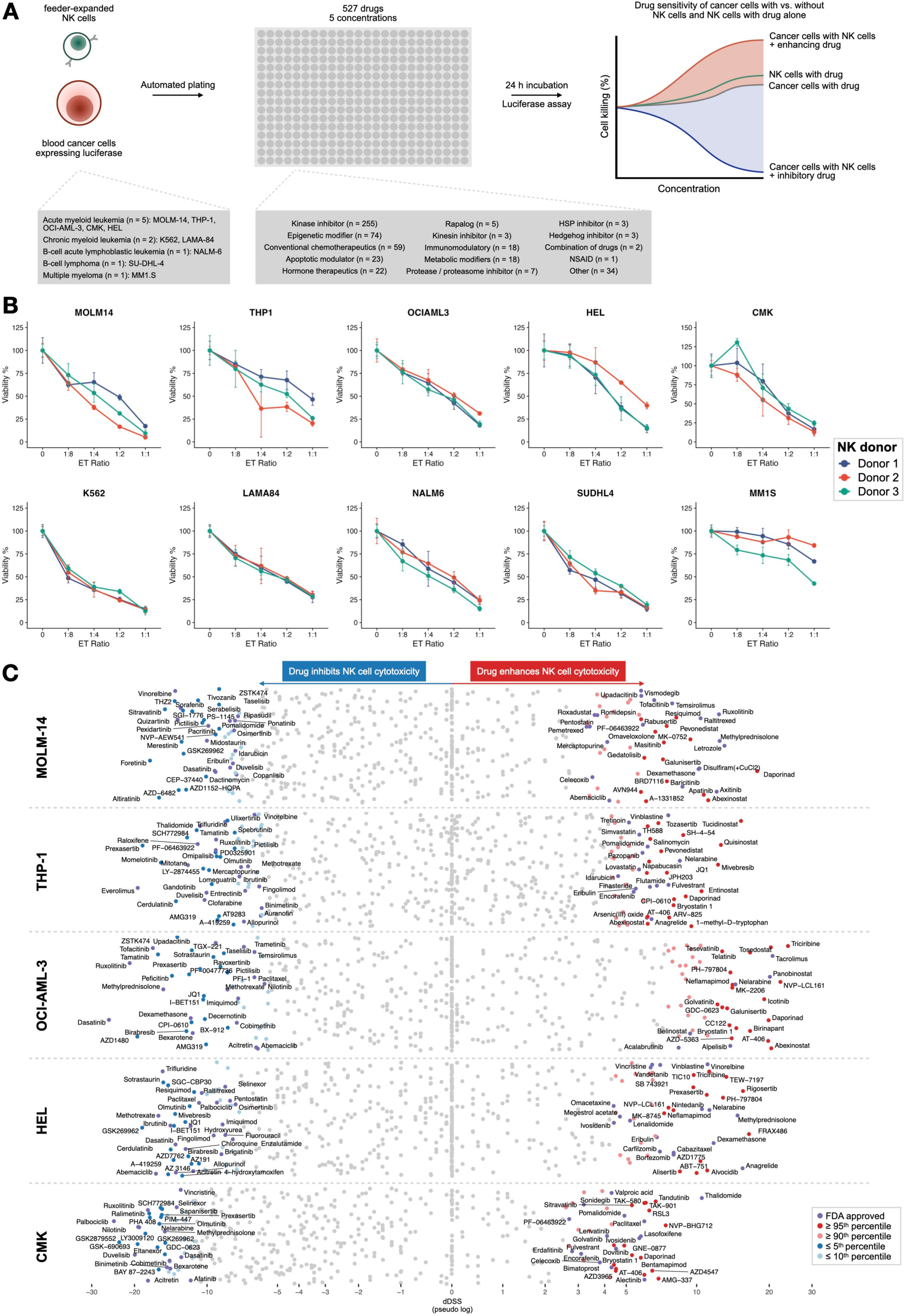
High-throughput drug resistance and sensitivity screening (DSRT) to identify modulators of NK cell cytotoxicity. (A) Overview of the high-throughput drug sensitivity and resistance testing workflow of drug-NK cell combinations using the FO5A drug library. (B) ET ratio titration of different NK cell donors against cell lines used in the study to identify ratios leading to 50% cancer cell inhibition. (C) Overview of how different compounds affect NK function in AML cell lines: Average pseudo-log transformed dDSS results from 5 AML cell lines (MOLM-14, THP-1, OCI-AML-3, HEL, CMK).

**Supplemental Figure 2:**
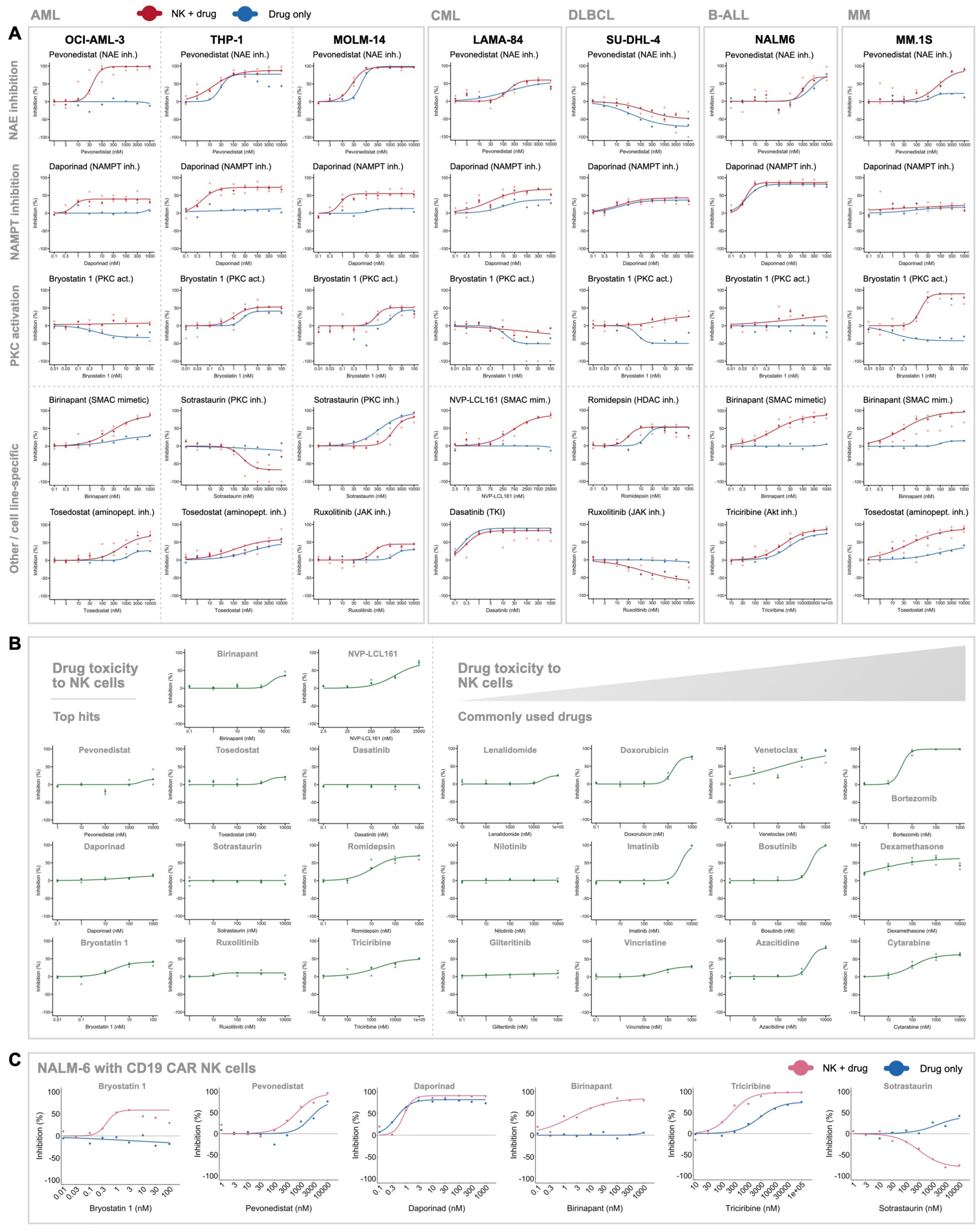
Impact of drugs on the cytotoxicity and viability of expanded NK cells and CAR NK cells. (A) Inhibition curves of top enhancers and inhibitors of NK cell cytotoxicity among cell lines representing different hematological malignancies. Each plot contains data from 3 replicate experiments using different NK cell donors. (B) Inhibition curves of NK cells upon treatment with drugs from the FO5A library, indicating drug toxicity to NK cells. (C) Inhibition curves from validated drugs in combination with CD19-targeting CAR NK cells, targeting NALM-6 cells.

**Supplemental Figure 3:**
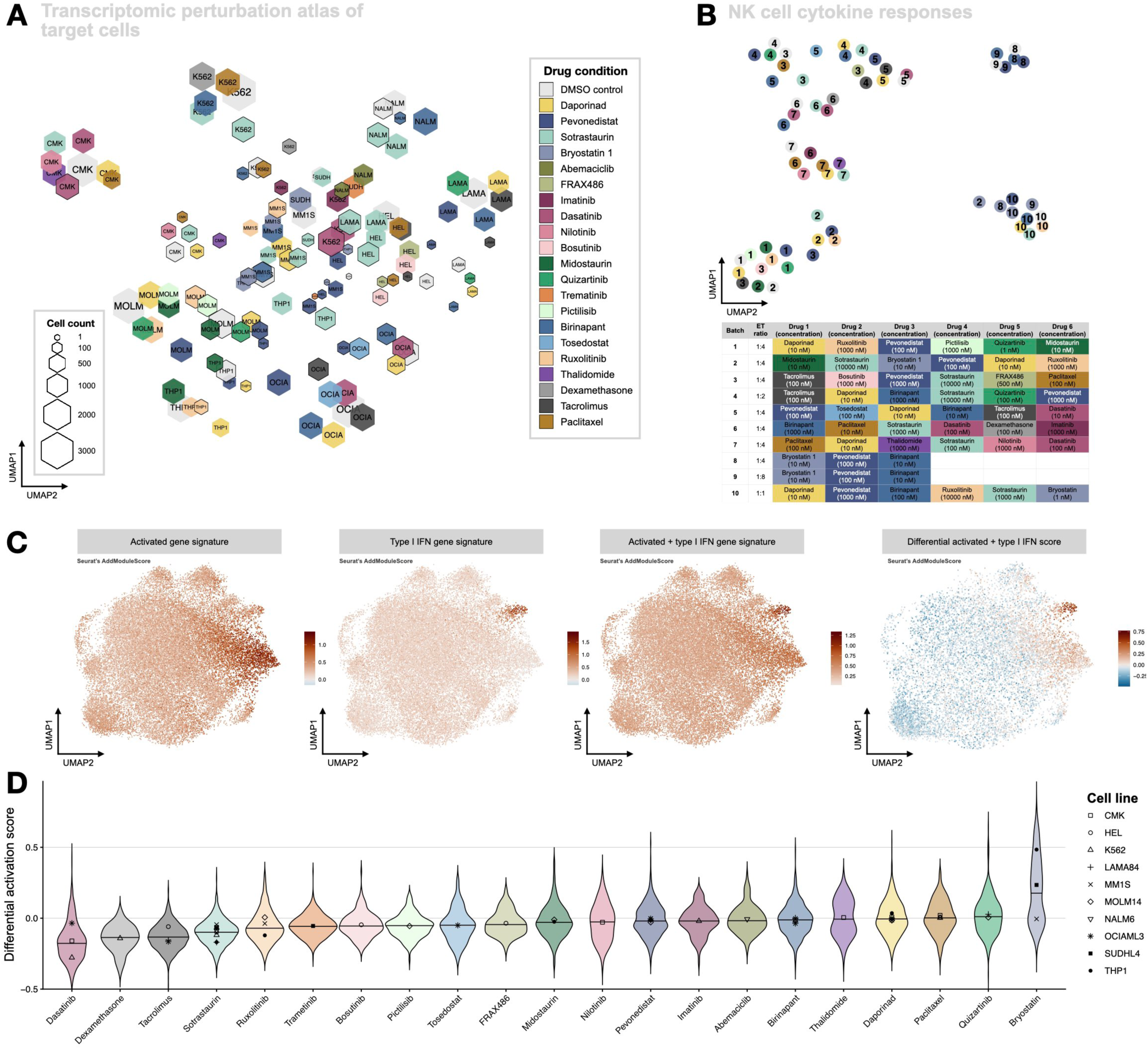
Overview of drug effects on NK and cancer cell transcriptome and cytokine secretion. (A) Overview of target cells’ scRNA-seq experiments as pseudo-bulk UMAP, showing varying transcriptomic changes depending on culture conditions and drug treatment. Symbols without and with borders represent mono- and cocultured conditions, respectively. (B) Overview of NK mono-cultured NK cells’ cytokine secretion depending on drug treatment and batch, visualized as UMAP of z-score-scaled cytokine levels. (C) Scores calculated using Seurat’s AddModuleScore function, reflecting the expression of previously published gene signatures^32^ corresponding to activated, type I IFN and a combination of the two, visualized as UMAPs. The rightmost UMAP shows the differential score of activated and type I IFN gene signatures. (D) Drug treatments’ effects on NK activation, visualized as violin plots of NK cell differential activation scores of drug-treated vs non-treated co-cultured NK cells. The scores were computed as the difference between mean activation scores from drug- and DMSO-treated cocultured NK cells based on previously published genes^32^ found upregulated in activated and type I IFN clusters, as illustrated in Figure S3C. Horizontal lines indicate median differential scores. The target cell line used in each co-culture is indicated by symbols in the violin plot to highlight differences between responses to different target cell lines.

**Supplemental Figure 4:**
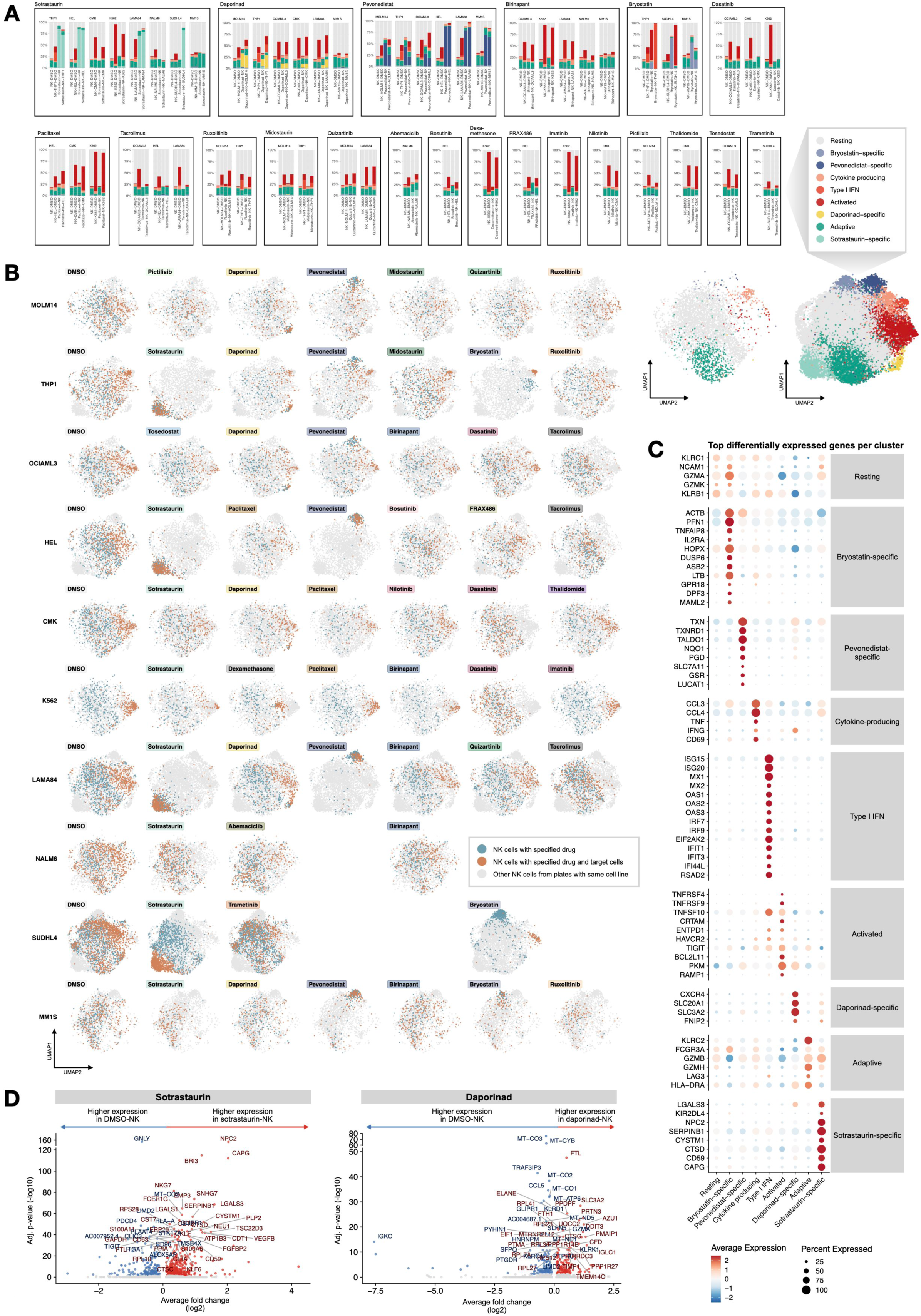
Evaluation of transcriptomic changes in NK cells induced drug treatment and co-culture. (A) Drug treatment-induced changes to NK cell phenotypes shown by a bar plot of percentages of NK cells per cluster by culture condition across all drug conditions tested using scRNA-seq. (B) UMAPs reflecting drug effects on NK cells in mono- and co-culture conditions, visualized separately for each drug tested in different cell lines. UMAPs of NK cells alone in DMSO and overall clustering of NK cells for reference visualized on the right. (C) Dot plot of top differentially expressed genes in NK cells for each cluster, selected based on relevance to NK cell function. (D) Sotrastaurin (left) and daporinad (right) -induced transcriptomic changes to NK cells compared to controls, shown as volcano plots of differentially expressed genes.

**Supplemental Figure 5:**
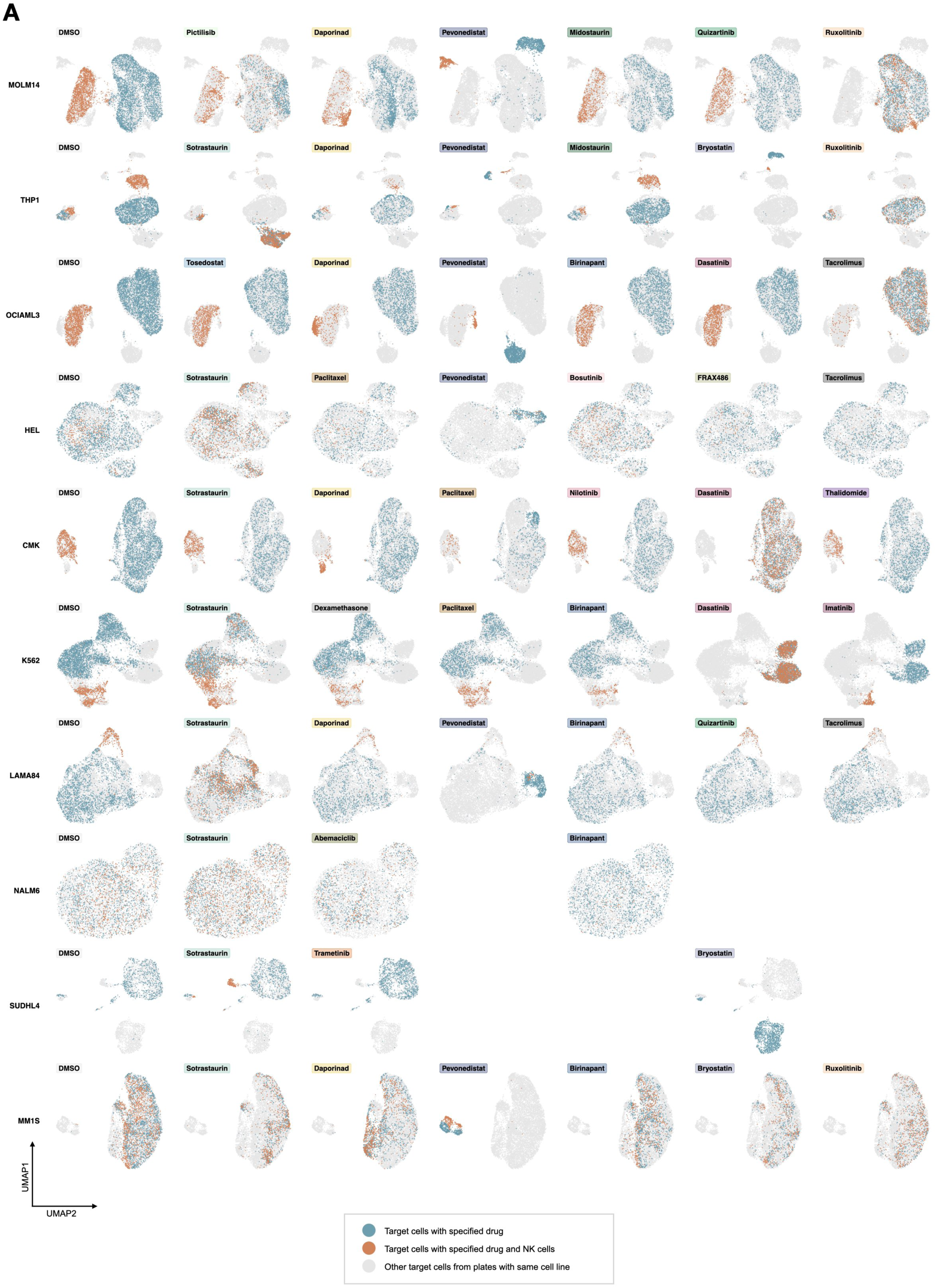
Evaluation of transcriptomic changes in cancer cells induced drug treatment and co-culture. (A) UMAPs reflecting drug effects on cancer cells in mono-and co-culture conditions, visualized separately for each drug tested in different cell lines. Colors reflect different culture conditions, with mono-culture colored in blue and co-culture conditions in orange.

**Supplemental Figure 6:**
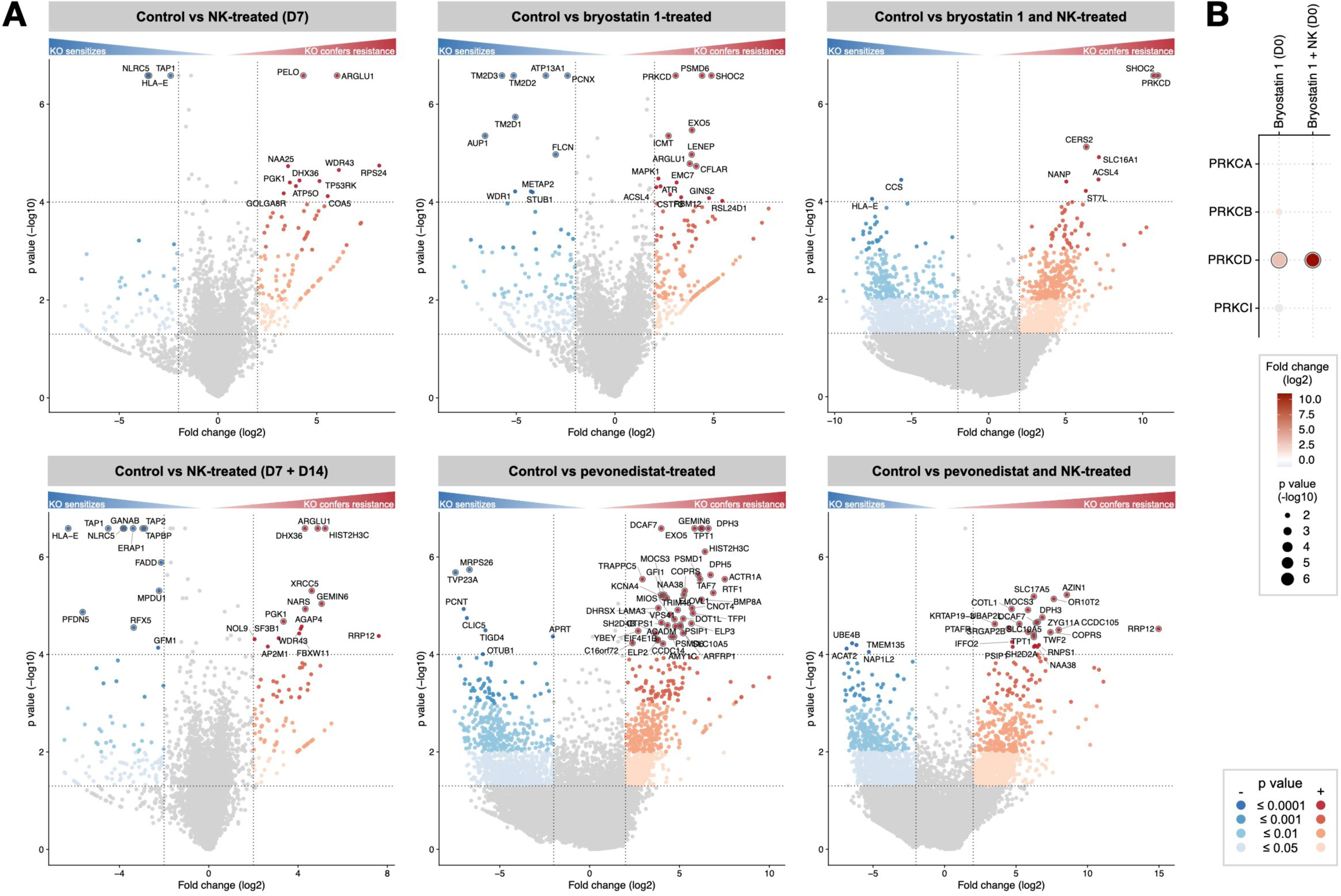
Genome-wide CRISPR screens of target cells treated with NK cells or drugs, or their combination. (A) Volcano plots of CRISPR results highlighting gene perturbations driving resistance or sensitivity to the combination of NK cells and bryostatin 1 (top) or pevonedistat (bottom). Treated conditions are compared to untreated controls, to identify treatment-specific gene perturbations. Gray circles around gene points indicate a false discovery rate (fdr) ≤ 0.05. (B) Dot plot of gene perturbations of PKC isoforms, with p-value ≤ 0.05, log2 fold change ≥ 1, fdr ≤ 0.7. The dark circle indicates fdr ≤ 0.05.

**Supplemental Figure 7:**
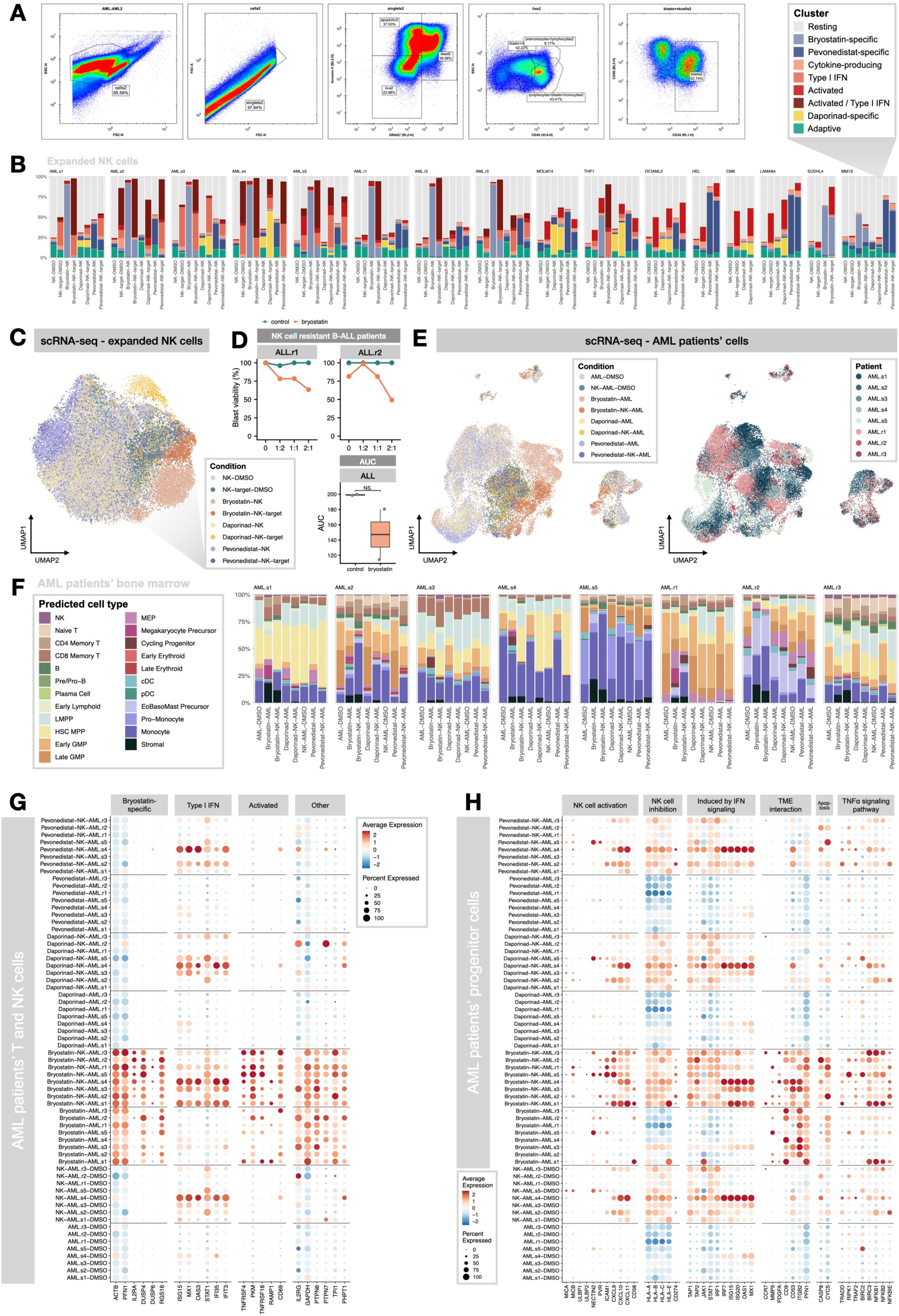
Validation experiments using AML and ALL patient samples. (A) Flow cytometry gating strategy for AML bone marrow samples treated with expanded NK cells and/or selected drugs, with the aim of gating leukemic blasts. (B) Drug treatment-induced changes to NK cell phenotypes shown by a bar plot of percentages of NK cells per cluster by culture condition across all drug conditions tested using scRNA-seq. Bar plots on the left reflect NK cell phenotypic distribution in AML patient sample experiments, and rightmost plots show distributions in cell line experiments. (C) UMAP visualization reflecting the distribution of NK cells within the UMAP based on treatment condition. (D) Flow-based killing assay readouts of ALL samples (top) and box plot of area-under-the-curve (AUC) values from ALL patient sample validation experiments. (E) UMAPs of AML patients’ cells, colored based on condition (left) and per patient (right). (F) Patient-wise bar plots indicating the distribution of patients’ bone marrow cells within BoneMarrowMap-predicted phenotypes. (G) Dot plots for individual conditions and patients, reflecting the expression of selected markers in AML patients’ NK and T cells (H) Dot plots for individual conditions and patients, showing the expression of genes affecting NK cell function in AML patients’ progenitor cells.

## DECLARATION OF GENERATIVE AI AND AI-ASSISTED TECHNOLOGIES IN THE MANUSCRIPT PREPARATION PROCESS

During the preparation of this work the author(s) used an AI-based language model (ChatGPT 5.0, OpenAI) to improve the clarity and style of the text. After using this tool/service, the author(s) reviewed and edited the content as needed and take(s) full responsibility for the content of the published article.

## Notes

### Competing Interest Statement

The authors have declared no competing interest.

